# Programmable Aptamer-Embedded Circular RNAs for Targeted Antigen-Presenting Cells Immunotherapy

**DOI:** 10.1101/2025.09.28.679023

**Authors:** Yu Zhang, Wenxi Wang, Xu Qian, Jian Chen, Ding Ding, Shaoju Gan, Weicheng Yi, Yalin Li, Yaping Deng, Zhongchao Chen, Hongyu Wu, Kedan Gu, Yuji Niu, Juanjuan Zhou, Xiangxin Kong, Qing Huang, Mengke Su, Luting Hu, Yingying Zhu, Meng Wang, Yuping Zhu, Shiyu Yao, Chao Zuo, Jing Zhang, Zhen Du, Liang Zhang, Xiaoman Zhai, Ting Fu, Yanjun Zhang, Jianmin Jiang, Yupeng Feng, Penghui Zhang, Xiangsheng Liu, Zhengbo Song, Sitao Xie, Weihong Tan

## Abstract

The targeted delivery of mRNA to specific cell types *in vivo* remains a major unmet need in mRNA-based therapeutics. Conventional strategies rely on nanocarriers engineered with various modifications. However, these exogenous materials could cause adverse reactions, and their complex manufacturing processes often hinder clinical application. By leveraging the innate programmability of RNA sequences, we herein report a carrier-free, scalable, and programmable aptamer-embedded circular RNAs (Apt-circRNAs) design that confers targeting capability through the strategic incorporation of different aptamers at varying densities, while retaining circular RNA stability and protein production capability. Experimental results show that antigen-loaded Apt-circRNAs enable targeted delivery to antigen-presenting cells (APCs), drive endogenous antigen expression, activate antigen-specific T cells, and elicit potent immune responses that mediate clearance of both early-and late-stage tumors in mouse models. In humanized models of colorectal cancer, the combination of Apt-circRNA and PD-1 blockade produced synergistic antitumor effects and significantly reshaped the tumor microenvironment. In a first-in-human (FIH) clinical trial, Apt-circRNA vaccine encoding KRAS G12D/G12V neoantigens exhibited a favorable safety profile and induced robust immune activation, T cell mobilization, and neoantigen-specific immune responses. With this study, we have established a new paradigm in carrier-free, APC-targeted RNA vaccine design, offering a safe and effective platform for next-generation cancer immunotherapy.

## Introduction

mRNA therapies are rapidly emerging as a new class of drugs with the potential to treat a wide range of diseases, including cancer^(1–8)^, autoimmune disorders^(9)^, and hereditary conditions^(10)^. The efficacy of mRNA drugs depends on their ability to deliver mRNA efficiently to specific cell types. Consequently, substantial research efforts are underway to develop engineered nanomaterial-based delivery vehicles for targeted and efficient mRNA therapeutics^(11–16)^. However, the introduction of exogenous delivery vectors often increases the complexity of manufacturing and raises concerns about potential unintended side effects *in vivo*, thus impeding clinical translation. Without requiring an external delivery carrier, naked mRNA can be directly delivered into cells through various techniques, including *in situ* electroporation, intradermal/intranodal/intratumoral injection, ultrasound-guided percutaneous injection, or jet injection^(17–25)^, enabling the expression of designed proteins and inducing antigen-specific humoral or cellular immune responses. However, unprotected mRNA is highly susceptible to rapid nuclease degradation *in vivo* and exhibits no inherent tropism for target cells, substantially restricting its biomedical utility^(26)^.

GalNAc-ligand based technology, rather than nano delivery, is now a clinically validated and effective method for targeted siRNA delivery to hepatocytes^(27–29)^. Moreover, scientists are investigating various targeted ligands (e.g., lipids, antibodies, aptamers) to extend siRNA therapeutics to extrahepatic targets^(30, 31)^. Our previous work established aptamer-mediated delivery as an efficient strategy for functional nucleic acid transport across biological barriers, including blood-brain barrier traversal to inhibit tau phosphorylation and thereby mitigate neuropathology.^(32)^. Moreover, aptamer-based systems have shown efficacy for delivering antisense oligonucleotides (ASOs), enabling precise *in vivo* targeting with reduced dosage, minimized off-target effects, and enhanced safety^(33–38)^.

Prompted by the success of targeted ligands in enabling precise RNA drug delivery, we propose the development of programmable aptamer-embedded circular RNAs (Apt-circRNAs) encoding tumor-specific antigens to target antigen-presenting cells *via* aptamer-mediated binding for cancer immunotherapy. The Apt-circRNA architecture comprises three functional modules. First, a targeting module consists of a rationally designed antigen-presenting cell (APC)-specific aptamers embedded at specific sites within the circular RNA scaffold. This preserves molecular stability and translational efficiency, while conferring high-affinity target recognition. Second, a stable expression framework, which is built on an engineered self-splicing circular backbone with low immunogenicity and superior nuclease resistance, enables prolonged structural integrity and sustained antigen expression *in vivo*^(39)^. Third, an antigen-encoding region features optimized non-repetitive Internal Ribosome Entry Site (IRES) elements and codons to drive robust expression of MHC-I/II-restricted antigens. This module allows flexible incorporation of diverse tumor neoantigens, viral antigens, or universal tumor-associated antigens, supporting rapid and modular vaccine development.

Apt-circRNAs were produced *via* an integrated *in vitro* transcription and circularization strategy, circumventing the limitations (e.g., inconsistent particle size or surface properties) of conventional nano delivery or ligand-based delivery systems and simultaneously guaranteeing high scalability and reproducibility. Upon subcutaneous administration, Apt-circRNAs are efficiently internalized by local APCs through aptamer-mediated targeting and then traffic to draining lymph nodes (dNLs). Apt-circRNA vaccines encoding mono-or multivalent antigens significantly suppressed tumor progression in syngeneic models, including poorly immunogenic types, both as standalone therapies and combined with immune checkpoint blockade (ICB). In a first-in-human (FIH) studies, the Apt-circRNA vaccine demonstrated an excellent safety profile; only one transient flu-like event (1/12) was reported, and no injection-site reactions or grade ≥2 adverse events occurred (0/12). The vaccine rapidly induced robust innate and adaptive immunity with response peaks returning to baseline shortly after each dose, indicating a transient, yet reproducible, immune cycle with controllable antigen-specific activation. These results underscore the clinical potential of Apt-circRNA, combining targeted immunogenicity with a favorable safety profile.

## Results

### Design and optimization of Apt-circRNAs for precise APCs targeting

Apt-circRNAs comprise three essential components: (1) a codon-optimized, antigen-encoding sequence, (2) an internal ribosome entry site (IRES) for ribosome recruitment and translation initiation, and (3) dendritic cell (DC) surface protein-targeting aptamers, facilitating targeted delivery to antigen-presenting cells (APCs) *in vivo*. Using an optimized permuted intron-exon (PIE)-based ribozyme system, we enable the efficient, scalable, and high-yield synthesis of Apt-circRNAs free of extraneous sequences^(40–43)^. We systematically engineered both the Anabaena pre-tRNA (Ana) and T4 bacteriophage td (T4td) intron-derived PIE platforms by replacing their extraneous fragments with engineered aptamers (Fig. 1a and Fig. S1). Our design capitalizes on the aptamer’s stem-loop structure by incorporating a cleavage site within the loop region. Following cleavage, the resulting aptamer fragments are ligated to a group I intron. To achieve complementarity with sequences flanking the aptamer cleavage site, we introduced mutations specifically into the intron’s P1 and P10 guide sequences. These components, along with other functional sequences, were then assembled into a recombinant RNA construct devoid of redundant elements. Critically, our strategy preserves the intact aptamer sequence. Instead of mutating the aptamer itself, we selectively engineered the guide sequences of group I intron to complement the sequence within the aptamer’s loop. This ensures precise, ribozyme-catalyzed splicing exclusively at the predefined loop site, generates Apt-circRNA products free of residual intron sequences, and eliminates the requirement for mutagenizing the aptamer to conform to splice site constraints. Specifically, the stem of the aptamer’s inherent stem-loop structure allows the ribozyme to generate an independent splicing bubble, which promotes efficient circularization of Apt-circRNAs. Successful circularization was verified by Sanger sequencing and gel electrophoresis (Fig. 1b,c and Fig. S1c,d), yielding an estimated circularization rate exceeding 80% by the Ana and T4td intron (Fig. S1e).

**Fig. 1.**
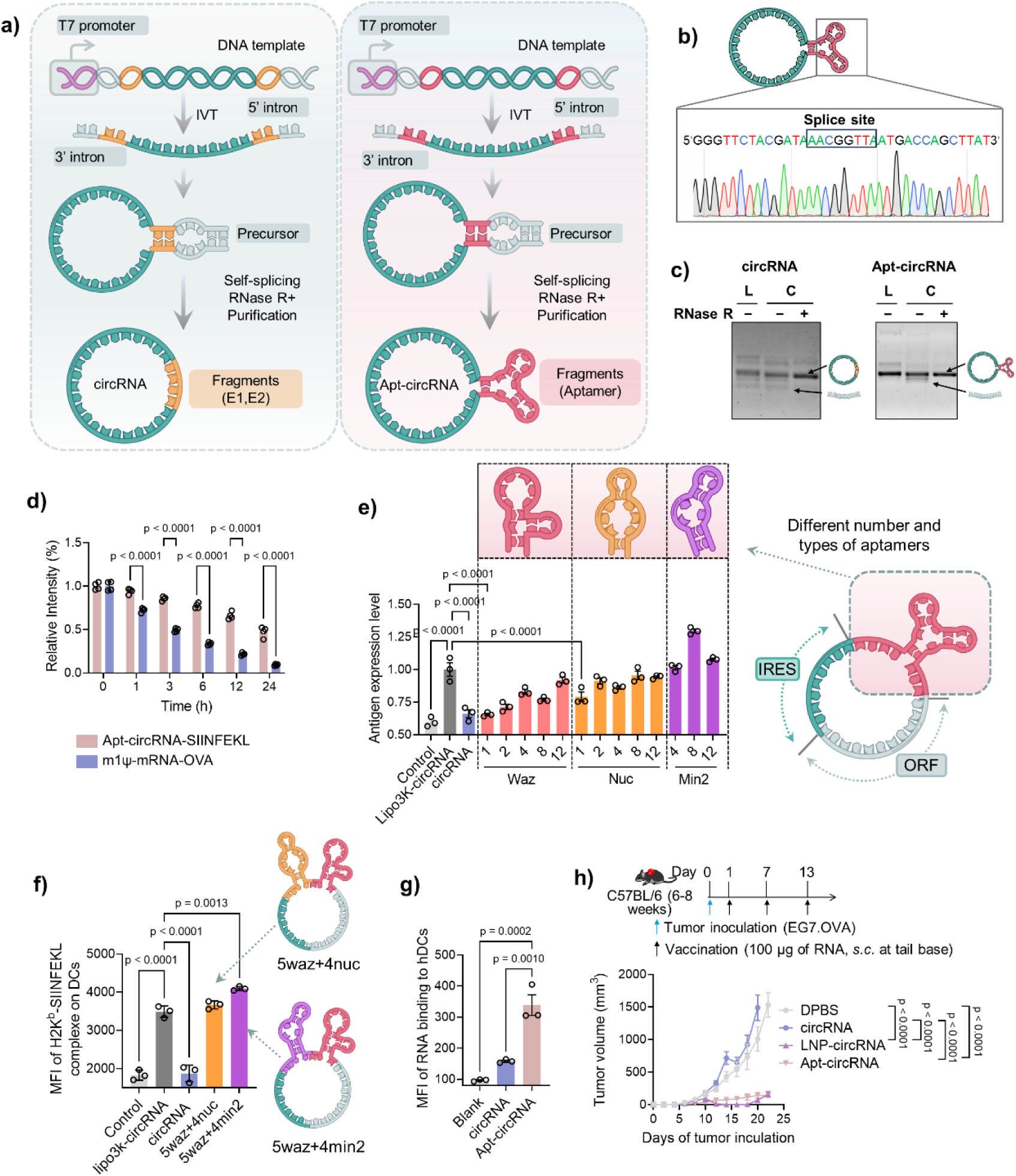
Design and optimization of Apt-circRNAs for precise APCs targeting. a, Schematic illustration of the circularization strategy based on PIE. b, Sanger sequencing of cDNA derived from Apt-circRNA indicates precise and uniform ligation of RNA precursors into circRNA by the T4 bacteriophage td (T4td) intron-derived PIE system. The denoted RNA sequence is converted from the Sanger sequencing results of cDNA. c, Circularized RNA products were analyzed with a denaturing agarose gel. Bands of circRNAs were verified with RNase R. d, Intact circRNA percentages of Apt-circRNA-SIINFEKL (0.3 mg/mL) and m1Ψ-mRNA-OVA (0.3 mg/mL) after incubation for different times in 10% FBS in PBS (37 °C). Data were quantified from gel electrophoresis by normalizing the band densities of different times to that of 0 hour (t-test). Data represent mean ± s.e.m, n=4. e, Antigen expression levels derived from SIINFEKL antigen presentation on BMDC cells treated with Apt-circRNA with different number and types of aptamers. Levels were normalized to those measured in BMDCs transfected with circRNA-SIINFEKL using Lipofectamine 3000 (Lipo3K-circRNA). Data represent mean ± s.e.m., n=3. f, Flow cytometric quantification of the mean fluorescence intensity (MFI) of SIINFEKL/H-2K^b^ complexes on BMDC cells treated with Apt-circRNA containing a combination of 5 waz aptamer and 4 min2 aptamer, as well as a combination of 5 waz aptamer and 4 nuc aptamer, LNP-circRNA-SIINFEKL, and controls, respectively, for 24 h. Data represent mean ± s.e.m., n=3. g, Apt-circRNA strongly binds human DCs. Data represent mean ± s.e.m., n=3. h, Top: design of EG7.OVA tumor immunotherapy studies in mice. Tumor cells were inoculated s.c. in mouse flank, and treatment started at day 1. Vaccines (circRNA: 100 μg) were administered by s.c. injection at the tail base. Bottom: Average tumor volumes after treatment with Apt-circRNA, LNP-circRNA, and free circRNA (n = 7). Data represent mean ± s.e.m. *P < 0.05, **P < 0.01, ***P < 0.001, ****P < 0.0001, one-way ANOVA with Bonferroni post-test.

The mRNA could be degraded by ubiquitous biological and environmental RNases, especially mRNA-degrading exonucleases, thereby compromising its shelf-life and biological half-life and, hence, limiting the efficiency and duration of antigen translation and, finally, immunomodulatory efficacy. However, in our construct, the absence of free 5’/3’ termini in circular RNAs confers intrinsic resistance to exonuclease-mediated degradation, resulting in much more biostability compared to linear mRNAs^(39, 44)^. To assess the stability of Apt-circRNAs, we employed commercially available linear mRNA modified with N1-methylpseudouridine (m1Ψ) as a control (m1Ψ-mRNA). Gel electrophoresis analysis revealed that Apt-circRNAs exhibited superior stability compared to that of the control mRNA, maintaining structural integrity in 10% FBS for over 24 hours (Fig. 1d and Fig. S2a). Since Apt-circRNA undergoes endosomal transport processes in different pH environments when entering the cell, we evaluated the stability of Apt-circRNAs under varying pH conditions. To accomplish this, Apt-circRNAs were incubated in buffers at pH 4.0, pH 7.2, and pH 8.0 for different durations at 37°C, followed by analysis using denaturing formaldehyde-agarose gel electrophoresis (Fig. S2b-c), and the results demonstrated that Apt-circRNAs remained stable within the pH range of 4.0 to 8.0 for over 24 hours.

For DC-targeted delivery, we selected three RNA aptamers specific for the following DC surface markers: nucleolin (nuc)^(45)^, transferrin receptor (waz)^(46, 47)^, and DEC-205 (CD205, an endocytic receptor; min2)^(48)^. Binding analysis using FITC-labeled aptamers revealed that waz exhibited superior affinity for both murine CD11c^+^ bone marrow-derived dendritic cells (BMDCs) and human DC populations compared to the other candidates (Fig. S3a-c). We then optimized the number and combination of aptamers incorporated into the circRNA scaffold. Regardless of the type of aptamer used, we observed that the expression of surface antigens on DCs gradually increased with higher aptamer quantities and that the min2 aptamer demonstrated enhanced antigen presentation (Fig. 1e). Building on evidence that bispecific aptamers enhance specific binding to target cells^(36, 49)^, we accordingly designed constructs combining two aptamer types. Screening assays identified an Apt-circRNA construct incorporating five waz and four min2 aptamers as inducing the optimal antigen presentation (Fig. 1f). Consequently, this combination was used for subsequent Apt-circRNA constructs.

Consistent with its design, Apt-circRNA demonstrated significantly increased binding to both murine BMDCs and human DCs compared to non-aptamer-bearing circRNA (Fig. 1g and Fig. S4). Furthermore, Apt-circRNA enabled rapid cellular uptake with a detectable Apt-circRNA fluorescence signal occurring within 4 hours post-transfection in cultured DCs (Fig. S5). Also, Apt-circRNAs encoding different lengths of ovalbumin (OVA) antigens (OVA_257-264_ (SIINFEKL, 8aa), OVA_230-264_ (35aa), OVA_257-386_ (130aa), and full-length OVA protein (386aa)) could also be presented on the surface of DCs (Fig. S6a).

To evaluate *in vivo* immunogenicity, we utilized the MHC class I-restricted ovalbumin epitope OVA_257-264_ (SIINFEKL) as a model antigen. Following subcutaneous injection at the tail base of C57BL/6 mice, H-2K^b^/SIINFEKL tetramer staining revealed that Apt-circRNA induced a frequency of SIINFEKL-specific CD8^+^ T cells in peripheral blood mononuclear cells (PBMCs) comparable to that induced by LNP-delivered circRNA, using the ionizable lipid SM-102 (Fig. S6b-d). We further assessed the antitumor efficacy of Apt-circRNA vaccines in a murine subcutaneous EG7.OVA tumor model. Apt-circRNA vaccine exhibited tumor growth inhibition comparable to that of LNP-circRNA, while non-aptamer-bearing circRNA showed no significant inhibitory effect (Fig. 1h and Fig. S6e,f).

### Apt-circRNAs efficiently targeted lymph nodes and APCs

To investigate the metabolic profile and biodistribution kinetics of Apt-circRNAs *in vivo*, we performed *in vivo* tracking using positron emission tomography (PET) imaging. Specifically, we radiolabeled Apt-circRNAs with ⁸⁹Zr by conjugating desferrioxamine (DFO) to a peptide nucleic acid (PNA) complementary to Apt-circRNAs. The radiolabeled Apt-circRNAs were administered *via* subcutaneous injection at the tail base of mice, followed by PET imaging over 24 hours. A circRNA without the aptamer was used as the control. By quantitative analysis of decay-corrected PET data, we found only minimal accumulation of control circRNA in the inguinal (IN) and axillary (AX) lymph nodes (LNs) with tissue uptake values below 0.1% of the injected dose per gram (%ID/g). In contrast, Apt-circRNAs accumulated in IN LNs within 0.5 hours post-injection. It subsequently drained to AX LNs by 12 hours and persisted within the LNs throughout the 24-hour imaging period (Fig. 2a,b). At 24 hours, Apt-circRNAs uptake reached 15.7 ± 2.6 %ID/g in IN LNs and 14.8 ± 0.3 %ID/g in AX LNs. These values significantly exceeded the uptake of unmodified circRNA (5.5 ± 2.8 %ID/g in IN LNs and 4.3 ± 2.9 %ID/g in AX LNs) (Fig. 2c). PET imaging showed predominant renal accumulation of Apt-circRNAs, consistent with renal clearance. No notable accumulation was observed in other major organs, indicating high targeting specificity, minimal off-target effects, and favorable safety profile (Fig. 2c). Efficient LN delivery of Apt-circRNAs was further confirmed using near-infrared (NIR) Cy7-labeled method, which exhibited strong fluorescence in draining LNs (Fig. S7a,b).

**Fig. 2.**
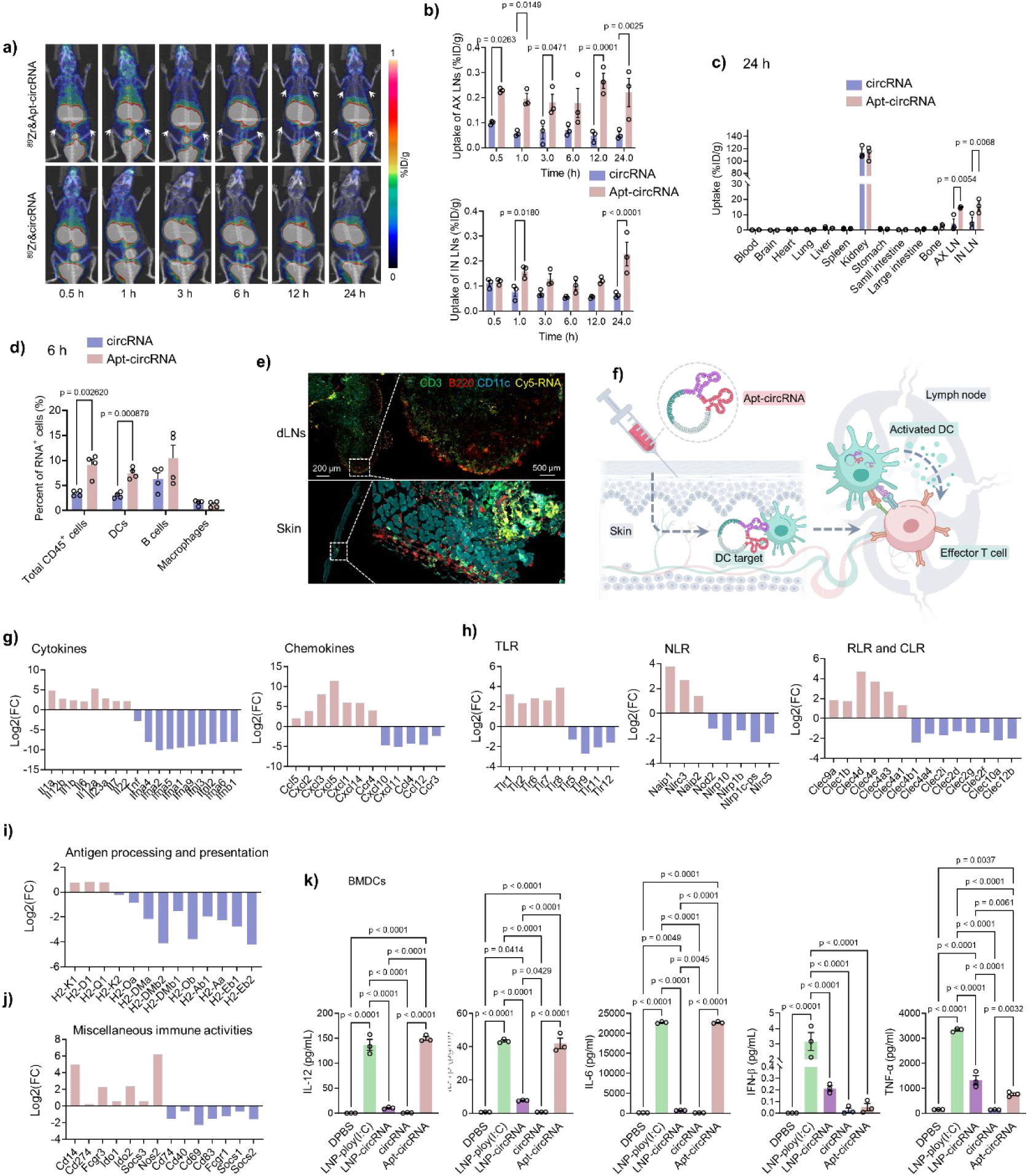
Apt-circRNA efficiently targeted lymph nodes and APCs, eliciting intrinsic innate immunostimulation. a, Representative coronal PET images showing C57BL/6 mice at different time post-injection of Apt-circRNA and circRNA at tail base (RNA: 0.5 nmol). b, Amount of Apt-circRNA and circRNA in IN and AX LNs quantified from three-dimensional (3D)-reconstructed, decay-corrected PET images. Data represent mean ± s.e.m., n=3. c, Biodistribution of tested compounds in resected organs measured by gamma counting at 24 h post-injection. Data represent mean ± s.e.m., n=3. d, Flow cytometry results showing Cy5-circRNA^+^ APC subsets among total CD45^+^ cells in draining lymph nodes 6 h post-injection of free Cy5-circRNA or Cy5-Apt-circRNA, respectively. Data represent mean ± s.e.m., n=4. e, Immunohistochemistry of skin and inguinal LNs 6 h after injection (CD3, green; B220, red; CD11c, blue; Cy5-Apt-circRNA, yellow). f, Schematic diagram of Apt-circRNA after subcutaneous injection in the body. g-j, Gene transcriptome analysis results from BMDCs transfected with Apt-circRNA, LNP-circRNA and PBS, respectively (24 h). Log_2_-transformed fold change (FC) represents log_2_ (ratio of mean expression induced by vaccine relative to PBS) (n = 3). Log_2_ (FC) of specific genes of interest related to cytokine and chemokine pathways (g), pattern-recognition receptor (PRR) pathways (h), antigen processing and presentation pathway (i), and miscellaneous immune activity pathways (j). k, Cytokines of BMDCs upon treatment with Apt-circRNA-SIINFEKL (10 μg/mL, 24 h) and controls. Poly(I:C) served as a positive control. Data represent mean ± s.e.m., n = 3. *P < 0.05, **P < 0.01, ***P < 0.001, ****P < 0.0001, one-way ANOVA with Bonferroni post-test.

We next analyzed the intranodal cell distribution of Cy5-labeled Apt-circRNA 6 hours after subcutaneous injection at the tail base. Administration of unmodified circRNA yielded minimal signal in draining IN LNs, whereas Apt-circRNAs administration resulted in much higher signal intensity (Fig. S7c). Following LN isolation and single-cell suspension preparation, flow cytometry detected the circRNA-Cy5 signal (Fig. S8). Distribution kinetics within key antigen-presenting cell subsets, including CD11c⁺ dendritic cells (DCs), B220⁺ B cells, and F4/80⁺ macrophages, was assessed (Fig. 2d). Consistent with PET imaging results, Apt-circRNAs increased cellular antigen uptake, resulting in a higher frequency of Cy5⁺ cells among total CD45⁺ cells and DCs. To elucidate the molecular mechanism of Apt-circRNAs trafficking to draining LNs, we performed histological analysis. Most Apt-circRNAs were efficiently internalized by DCs at the injection site, followed by accumulation in the subcapsular sinus and interfollicular regions of draining LNs (Fig. 2e). In contrast, lipid nanoparticle-encapsulated circRNA (LNP-circRNA) primarily remained as intact nanoparticles near the injection site before being taken up by B cells and DCs within LNs. No circRNA fluorescence signal was detected at the subcutaneous injection site in the group without aptamer, and only minimal signal was observed within the lymph nodes (Fig. S9). These findings suggest that the aptamer facilitates specific DC recognition and internalization of Apt-circRNAs, enabling targeted delivery to draining LNs (Fig. 2f).

### The intrinsic innate immunostimulatory property of Apt-circRNAs

Various RNA species activate innate immunity through endosomal pattern recognition receptors, or PRRs, which include Toll-like receptors 3, 7 and 8 (TLR3/7/8) and cytosolic PRRs, which include retinoic acid-inducible gene I (RIG-I) and melanoma differentiation-associated protein 5 (MDA5)^(50)^. This response provides cytokines and co-stimulatory signals essential for adaptive immunomodulation. The mRNA vaccines exploit the inherent immunostimulatory properties of nanocarriers like lipid nanoparticles (LNPs), leveraging innate immunity to provide antigen-presenting cells (APCs) with the proinflammatory cytokines and co-stimulatory signals required for antigen presentation and T cell priming.

To investigate the immunomodulatory impact of Apt-circRNA vaccines, we performed RNA-seq transcriptomic analysis on mouse bone marrow-derived dendritic cells (BMDCs) transfected with either Apt-circRNA or LNP-circRNA for 24 hours, using PBS or blank LNPs as controls. We categorized approximately 5379 differentially expressed genes (DEGs) between the Apt-circRNA and LNP-circRNA groups (Fig. S10a). Apt-circRNA induced 2433 upregulated and 2946 downregulated genes relative to LNP-circRNA (Fig. S10b). Gene ontology (GO) analysis indicated that upregulated genes were primarily enriched in processes related to inflammation and immune response, including protein binding, positive regulation of gene expression, inflammatory response, and immune response. In contrast, downregulated genes showed enrichment for cytokine receptor binding, type I interferon receptor binding, membrane localization, and nucleoplasm (Fig. S10c).

Specifically, transcript levels of key cytokines/chemokines (Il-6, Il-1α/β, Il-12α/β, Cxcl3, Cxcl5; Fig. 2g) and PRRs (Tlr2/6, Tlr7/8, Naip1; Fig. 2h) were much higher in Apt-circRNA-treated cells compared to those observed in the LNP-circRNA-treated cells. These distinct PRR activation profiles between Apt-circRNA and LNP-circRNA likely arise from structural differences. Moreover, Apt-circRNA promoted DC maturation, as evidenced by the expanded expression of MHC-I genes (H2-K1, H2-D1, H2-Q1; Fig. 2i, Fig. S11), and induced a multifaceted proinflammatory shift characterized by upregulation of cd14, Fcgr3, Ido2, and Nos2 (Fig. 2j).

Validation of DC inflammation using cytokine secretion assays (with poly(I:C) as a positive control) confirmed that Apt-circRNA treatment elevated the secretion of proinflammatory cytokines IL-1β, IL-6, and IL-12 relative to circRNA, LNP-circRNA and control treatments, consistent with transcriptomic data (Fig. 2k). These results provide a proof of concept for immunomodulatory mechanisms of Apt-circRNA vaccines that involve such complex processes as antigen presentation, PRR signaling pathways, cytokine secretion, and pro-inflammatory response. Importantly, Luminex assays revealed that Apt-circRNA elicited lower systemic levels of reactogenicity-associated chemokines and cytokines than those elicited by LNP-circRNA, except IL-12 (Fig. S12a,b), indicating a favorable safety profile. Apt-circRNA also demonstrated reduced cytotoxicity in BMDCs compared to that induced by LNP-circRNA (Fig. S12c).

Correspondingly, mice administered high-dose LNP-circRNA exhibited weight loss, whereas only minimal weight changes occurred in the Apt-circRNA group (Fig. S12d). Collectively, these findings preliminarily demonstrate the immunomodulatory mechanism of Apt-circRNA vaccines by combining antigen production and activation of targeted immune cells.

### Multivalent Apt-circRNA vaccines for robust combination immunotherapy

Modular Apt-circRNA vaccines can be easily adapted to encode diverse peptide immunogens for broad therapeutic applications. Accordingly, we first constructed a multivalent Apt-circRNA vaccine encoding the MHC-I-restricted Adpgk (MC38 tumor cell-specific neoantigen), MHC-I-restricted epitopes HPV16 E7_49–57_ (for HPV-associated cancers), and the melanoma-associated antigens mouse TRP2_180–190_ and human gp100_23–33_ (Apt-circRNA-AETG, Fig. 3a). We found that the Apt-circRNA-AETG neoantigen vaccine inhibited the growth of Adpgk-positive MC38 tumor (Fig. 3b). We further evaluated the therapeutic potential of Apt-circRNA-AETG in the E7-positive TC-1 tumor model, and monotherapy with Apt-circRNA-AETG was also shown to significantly suppress tumor progression (Fig. 3c). In the poorly immunogenic B16F10 melanoma model, Apt-circRNA-AETG again demonstrated marked tumor growth inhibition (Fig. 3d). However, when we combined Apt-circRNA and anti-PD-1 antibody, the resultant synergism potently strengthened therapeutic efficacy and prolonged mouse survival (Fig. 3d and Fig. S13). Notably, this combination led to complete tumor regression without recurrence in two out of seven mice over a three-month observation period.

**Fig. 3.**
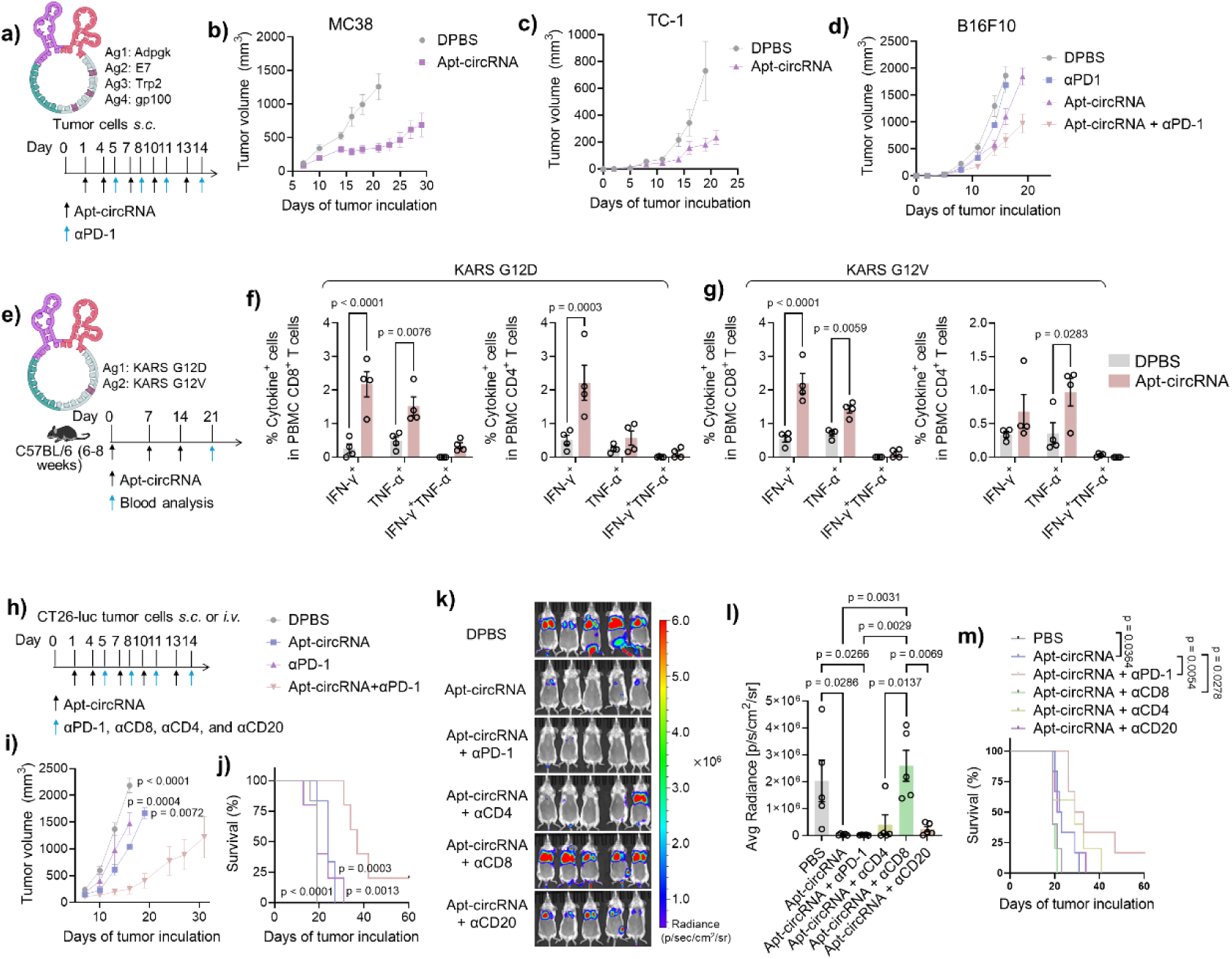
Multivalent Apt-circRNA vaccines for robust combination immunotherapy of multiple types of tumors. a, Top: schematic diagram of multivalent Apt-circRNA vaccine encoding the MHC-I-restricted Adpgk (MC38 tumor cell-specific neoantigen), MHC-I-restricted epitopes HPV16 E7_49–57_ (for HPV-associated cancers, such as TC-1), and the melanoma-associated antigens mouse TRP2_180–190_ and human gp100_23–33_ (Apt-circRNA-AETG). Bottom: design of MC38, TC-1, and B16F10 tumor immunotherapy studies in mice. Tumor cells were inoculated subcutaneously in mouse flank, and treatment started on day 1. Vaccines were subcutaneously injected at tail base; antibodies were intraperitoneally injected. b-c, Average MC38 (b) or TC-1 (c) tumor volumes after treatment with Apt-circRNA-AETG and controls. Data represent mean ± s.e.m., n=7. d, Average volumes of B16F10 melanoma treated with Apt-circRNA-AETG vaccine alone or combined with anti-PD-1. Data represent mean ± s.e.m., n=7. e, Top: schematic diagram of bivalent Apt-circRNA vaccine (Apt-circRNA-KR2) encoding G12D (G12D_2–23_) and G12V (G12V_2–23_) with mutations at residue 12. Bottom: timeline of vaccination and drawing blood. Vaccines were subcutaneously injected at tail base. f, Intracellular IFNγ and TNF staining of PBMC CD8^+^ and CD4^+^ T cells treated with G12D peptide from the above immunized mice. g, Intracellular IFNγ and TNF staining of PBMC CD8^+^ and CD4^+^ T cells treated with G12V peptide from the above immunized mice. Data represent mean ± s.e.m., n=5. h, Design of tumor immunotherapy studies in mice. Tumor cells were inoculated subcutaneously in mouse flank or intravenously, and treatment started on day 1. Vaccines were subcutaneously injected at tail base; antibodies were intraperitoneally injected. i, CT26 tumor growth after Apt-circRNA-KR2 vaccine treatment. Asterisks: statistical significance relative to Apt-circRNA-KR2 + anti-PD-1. Data represent mean ± s.e.m., n=7. j, Kaplan–Meier survival curves of as-treated CT26-bearing mice. Asterisks: statistical significance relative to Apt-circRNA-KR2 + anti-PD-1. k, *In vivo* imaging system (IVIS) images of CT 26-luc tumors on day 14. l, Fluorescence intensity of CT26-luc tumors in different groups on day 14. Data represent mean ± s.e.m., n=5. m, Kaplan-Meier survival curves of as-treated CT26-luc-bearing mice. *P < 0.05, **P < 0.01, ***P < 0.001, ****P < 0.0001, one-way ANOVA with Bonferroni post-test.

KRAS mutations lead to uncontrolled cell growth and are mainly observed in pancreatic cancer, colorectal cancer (CRC), and lung cancer with mutation frequencies of 97.7, 44.7, and 30.9%, respectively^(51)^. Current immunotherapy, such as immune checkpoint blockage (ICB) and adoptive cell transfer, is limited by ICB resistance and poor T cell memory. Since combinatorial immunotherapy administered *via* therapeutic vaccines could elicit a T cell response in KRAS-mutant cancer, a bivalent Apt-circRNA vaccine encoding the G12D (G12D_2–25_) and G12V (G12V_2–25_) with mutations at residue 12 (Apt-circRNA-KR2, Fig. 3e) was constructed accordingly. In Balb/c mice, Apt-circRNA-KR2 (100 μg; days 0, 7 and 14) elicited KRAS-specific polyfunctional CD8^+^ and CD4^+^ T cells, as shown by intracellular IFNγ/TNF (formerly known as TNFα) staining (day 21) (Fig. 3f,g and Fig. S14). Given the robust T cell response elicited by Apt-circRNA-KR2, we administered Apt-circRNA-KR2 to CT26 tumor-bearing mice with a KRAS G12D mutation to test its immunotherapeutic effects (Fig. 3h). As expected, compared with the DPBS control group, the Apt-circRNA-KR2 vaccine inhibited the growth of CT26 tumors (Fig. 3i). Moreover, combining the vaccine with anti-PD-1further potentiated the therapeutic efficacy (Fig. 3i) and prolonged mouse survival (Fig. 3j) with 2 out of 7 tumors showing complete regression without recurrence in 2 months.

We also evaluated the antitumor effect of Apt-circRNA-KR2 in the CT26-luc lung metastasis mouse model (Fig. 3h). Results showed that the Apt-circRNA-KR2 group exhibited fewer pulmonary metastatic nodules compared to the DPBS group, suggesting a more effective antitumor response from the Apt-circRNA-KR2 vaccines (Fig. 3k,l). To assess the relative significance of CD8^+^ versus CD4^+^ T cells and B cells in antitumor activities elicited by the Apt-circRNA-KR2 vaccine, *in vivo* anti-CD20, anti-CD4, and anti-CD8 antibodies were used to deplete B cells, CD4^+^ and CD8^+^ T cells, respectively (Fig. 3h). Notably, depletion of CD8^+^ T cells completely abolished the coinhibitory effects of Apt-circRNA-KR2, as supported by luciferase-based imaging analysis (Fig. 3k,l). In contrast, depletion of B cells and CD4^+^ T cells only minimally compromised the antitumor effects of Apt-circRNA-KR2 (Fig. 3k,l). At the 2-month follow-up, 1 of 6 mice in the Apt-circRNA-KR2 + anti-PD-1 groups achieved long-term disease-free survival, notably surpassing the survival rate of the DPBS group (0 of 6) and the Apt-circRNA-KR2 + anti-CD8 antibody group (0 of 6) (Fig. 3m). Collectively, our data suggest that CD8^+^ T cell-mediated immune response effectively contributes to tumor killing and growth control after Apt-circRNA-KR2 vaccine immunization.

NCG-M, a mouse model genetically modified to express human granulocyte– macrophage colony-stimulating factor and IL-3, was used to validate the clinical potential of our Apt-circRNA vaccines since it supports the engraftment of human myeloid and T cells. After transplanting human hematopoietic stem cells (HSCs) into irradiated NCG-M mice, those with around 20% huCD45^+^ cells in peripheral blood by week 7 were selected. By week 10 to week 14, a humanized patient-derived organoid xenograft (PDOX) model of microsatellite stable (MSS)/mismatch repair-proficient (pMMR) colorectal cancer (CRC) with fully differentiated macrophages and T cells had been established (Fig. 4a). The pathological sections of the tumor tissues from NSG mice were consistent with those from the patient’s tumor and the PDOX model, indicating that model construction was successful (Fig. 4b and Fig. S15). After the tumor model was successfully constructed, the Apt-circRNA-KR2 vaccine was injected into the mouse tail vein. The injection was given every three days for a total of six times. On the fifth day, anti-human-PD-1 antibody was injected intraperitoneally once every three days for a total of five times (Fig. 4c). Treatment with Apt-circRNA-KR2 significantly inhibited tumor growth in this humanized model (Fig. 4d). Moreover, when combined with the anti-PD-1 antibody, growth of the tumor was further inhibited (Fig. 4d). These findings underscore the effectiveness of Apt-circRNA-KR2 in a clinically relevant setting, highlighting its potential for future clinical applications.

**Fig. 4.**
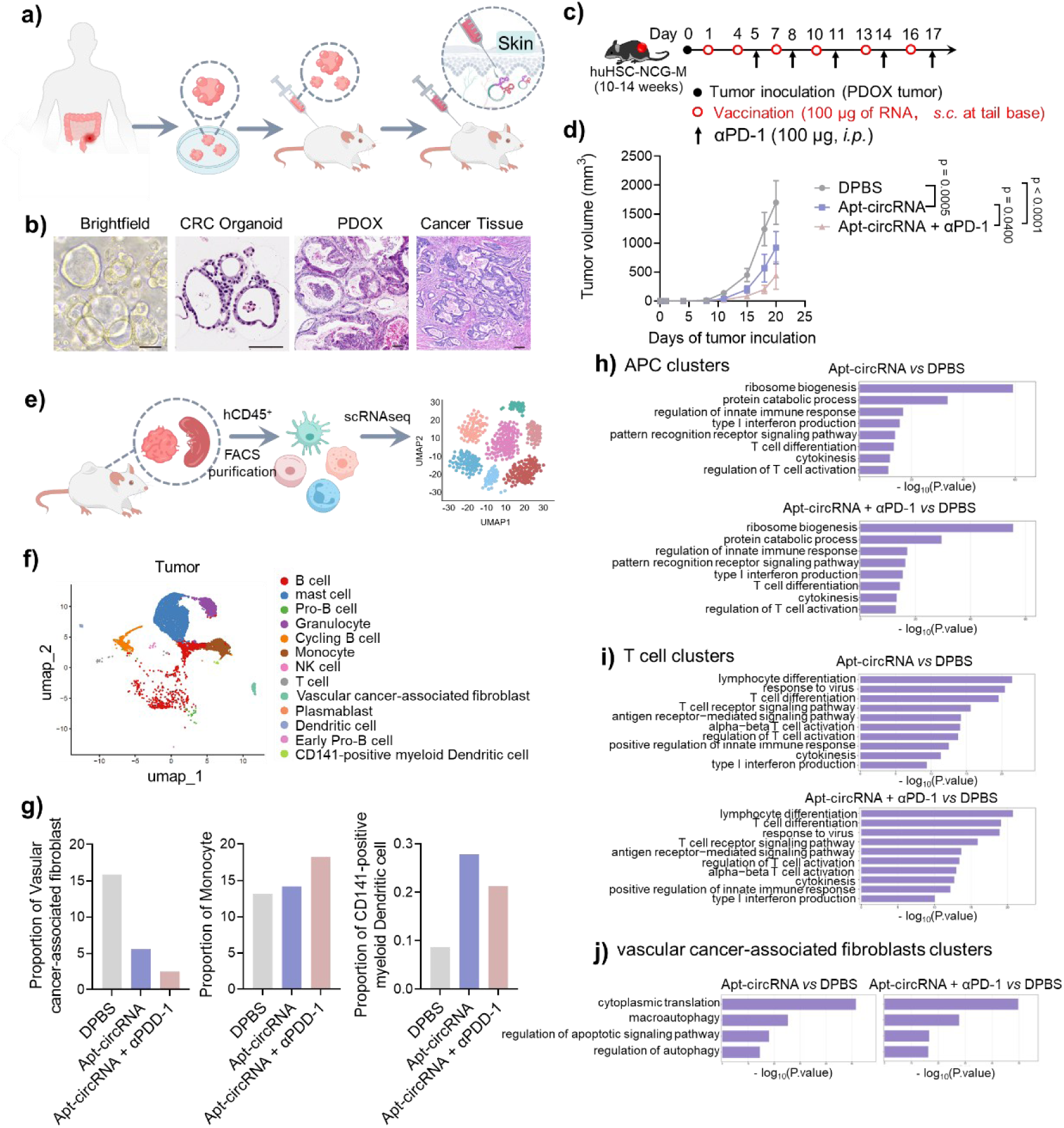
Therapeutic efficacy of Apt-circRNA in the hu-HSC-PDX CRC model. a, Schematic illustration of the experimental design for the hu-HSC-PDX-CRC model. b, H&E and IHC staining of Patient-017 tumor tissue, organoids derived from it, and the patient-derived organoid xenograft model (PDOX) designated as PDO-017. Organoids exhibit morphological features similar to those of the parent tissue. Scale bar, 100 µm. c, Timeline of vaccination and anti-PD-1 injection. Vaccines were subcutaneously injected at tail base; antibodies were intraperitoneally injected. d, Hu-HSC-PDOX-CRC tumor growth after Apt-circRNA-KR2 vaccine treatment. Data represent mean ± s.e.m., n=4. e, Study schema of scRNA-seq of tumor tissue and spleen in Hu-HSC-PDX-CRC on day 21. f, Uniform manifold approximation and projection (UMAP) representations of scRNA-seq data of all CD45^+^ cell subsets in tumor. g, Average proportion of vascular cancer-associated fibroblast, Monocyte, and CD141-positive myeloid dendritic cell type derived from the tumor tissues. h-i, Differential pathways enriched for the discriminative markers of APC (h) and T (i) cell subsets in tumor by GO terms. j, Differential pathways enriched for the discriminative markers of vascular cancer-associated fibroblast subsets in tumor by GO terms.

To explore the therapeutic mechanism of the Apt-circRNA vaccine in the NCG-M mouse model, we performed single-cell transcriptomic analyses (single-cell RNA sequencing (scRNA-seq)) of human CD45^+^ (hCD45^+^) cells in tumor tissues and spleen tissues (Fig. 4e). Ultimately, we identified 13 different cell types and states within the immune cell populations, including T cells, monocytes, mast cells, natural killer (NK) cells, B cells, and so on (Fig. 4f and Fig. S16a,b). In tumor tissues, compared with the control group, the proportion of vascular cancer-associated fibroblasts in both the Apt-circRNA and Apt-circRNA + anti-PD-1 antibody groups had been reduced, while the proportion of APCs, such as monocytes and CD141-positive myeloid dendritic cells (DCs), had increased (Fig. 4g). In spleen tissue, compared with the control group, the proportion of B cells in the vaccine group and the combined group decreased, while the proportion of monocytes increased (Fig. S16c). Among the APC clusters (monocytes, B cells, DCs and CD141-positive myeloid DCs), GO enrichment analysis pathways related to antigen translation and presentation, DCs maturation, and T cell differentiation induction and maintenance were significantly upregulated in the Apt-circRNA and combined groups (Fig. 4h and Fig. S16d), including ribosome biogenesis, tRNA processing, regulation of innate immune response, positive regulation of innate immune response, and T cell differentiation. For T cell clusters, GO enrichment analysis pathways related to proliferation of cytotoxic T cells were also significantly upregulated in both groups (Fig. 4i and Fig. S16e), including response to virus, T cell receptor signaling pathway, antigen receptor-mediated signaling pathway, alpha-beta T cell activation, cytokinesis, and type I interferon production. For vascular cancer-associated fibroblasts, GO enrichment analysis pathways related to apoptosis and autophagy were, again, significantly upregulated in both groups (Fig. 4j), including macroautophagy, regulation of apoptotic signaling pathway, and regulation of autophagy. In short, Apt-circRNA both expanded antigen presentation by efficiently expressing antigens in human APCs and remodeled the tumor immune microenvironment (TIME) by inducing autophagy and apoptosis in cancer-associated fibroblasts, thereby promoting cytotoxic T cell proliferation.

### Apt-circRNA-KR2 vaccination elicits potent immunity in human

The favorable preclinical outcomes observed in safety and efficacy studies using a GLP-grade Apt-circRNA-KR2 vaccine in rats (Fig. S17 and Fig. S18) and the NCG-M mouse model propelled momentum toward clinical translation. We therefore initiated a clinical trial at Zhejiang Xiaoshan Hospital to evaluate both the safety and early immune responses elicited by Apt-circRNA-KR2 in healthy volunteers.

First-in-human safety assessment of an Apt-circRNA vaccine enrolled 9 healthy volunteers (aged 18-55 years; Male ≥50 kg, female ≥45 kg; Body Mass Index (BMI): 19.0–26.0 kg/m² (inclusive)). Single dose-escalation (50, 100, and 250 μg; n=3/group) initiated at 50 μg (1/20 of rat maximum safe dose). Serial peripheral blood monitoring over 168 hours post-administration revealed that all hematologic parameters, immune cell subsets, and cytokines remained within normal ranges, demonstrating favorable tolerability across tested doses (Fig. S19-S21).

Following the single-dose safety profile, an expanded cohort of three HLA-A*02:01-or HLA-A*11:01-positive healthy volunteers (identical inclusion criteria) received multi-dose administrations across three escalating dose groups (250 μg) on days 1, 7, and 13 (Fig. 5a,b). The most common adverse event was a transient flu-like symptom in one participant, which resolved within 12 hours post-injection. No grade ≥2 (dose-limiting) toxicities were observed, and no skin inflammatory reactions occurred at the injection sites in any subject (Table 1 and Fig. 5b). Serial peripheral blood monitoring over 180 days post-administration revealed all hematologic parameters, immune cell subsets, and cytokines still remained within normal ranges, demonstrating favorable tolerability across tested doses and prolonged timespan (Fig. S22-S24). These results preliminarily demonstrated that Apt-circRNA-KR2 is well tolerated in humans.

**Fig. 5.**
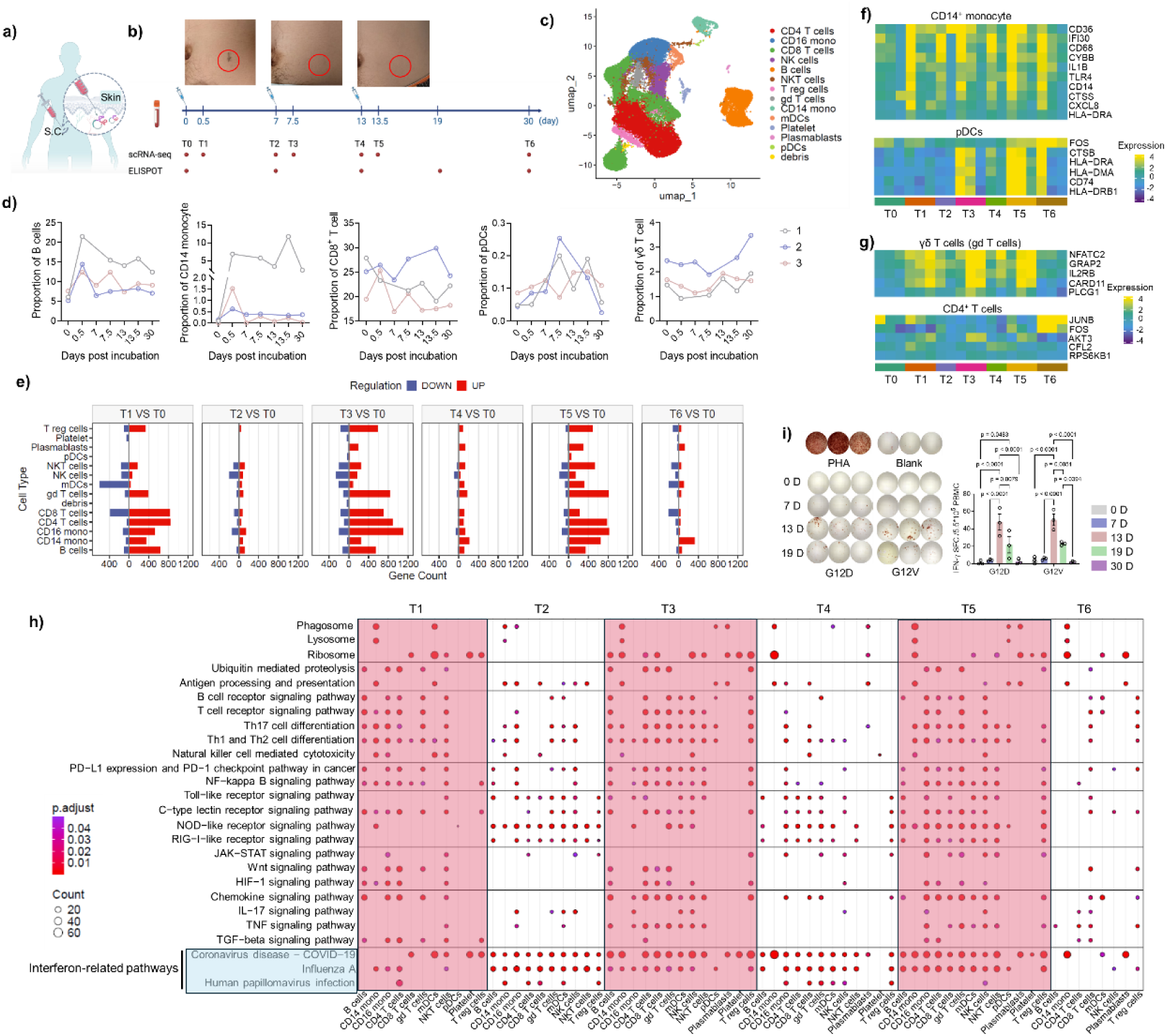
Single-cell transcriptional profiling of human PBMCs after Apt-circRNA vaccination. a, Overview of design and administration of Apt-circRNA vaccine. b, Top, example of an injection site reaction after vaccination. Bottom, Treatment and sample collection schedule. c, UMAP representation of scRNA-seq of PBMC populations (n = 3 volunteers). d, Average proportion of each cell type derived from human PBMCs after vaccination at different times. e, Heatmap showing the relative change in expression of total genes in lymphoid subsets before and after vaccination. f, Heatmap showing the gene expression of representative cell function-related genes in two major APC cell clusters. g, Heatmap showing the gene expression of representative cell function-related genes in two major T cell clusters. h, Highly enriched pathways among differentially expressed genes based on GSEA. i, IFN-γ ELISpot of serial post-vaccination PBMCs restimulated ex vivo with antigen peptide pools. Data represent mean ± s.e.m., n=3. *P < 0.05, **P < 0.01, ***P < 0.001, ****P < 0.0001, one-way ANOVA with Bonferroni post-test.

**Table 1:** Treatment-related adverse events.

| Events | Grade 1 | Grade 2-5 |
| --- | --- | --- |
|  | Number of healthy donors n<sup>a</sup> (%) |  |
| Upper respiratory tract infection | 1 (8.3) | 0 |
| Decreased basophil count | 2 (16.7) | 0 |
| Decreased lymphocyte count | 1 (8.3) | 0 |
| Decreased red blood cell count | 1 (8.3) | 0 |
| Hypotension | 1 (8.3) | 0 |
| Hypertriglyceridemia | 1 (8.3) | 0 |
| Increased monocyte count | 1 (8.3) | 0 |
| Interleukin 6 level increased | 4 (33.3) | 0 |
| Interleukin 10 level increased | 1 (8.3) | 0 |
| Interleukin 5 level increased | 1 (8.3) | 0 |
| Pyrexia | 0 | 0 |
| Liver function abnormality | 0 | 0 |
| Renal function abnormality | 0 | 0 |
| Pain | 0 | 0 |
| Injection site reaction | 0 | 0 |

<sup>a</sup>Number of healthy donors that experienced adverse events. Data are presented as a percentage of total donor cohort (n=12).

The observation of transient flu-like symptoms suggested that the vaccine might have triggered an early innate response. To characterize vaccine-induced immunity in greater detail, we performed scRNA-seq on PBMCs from three participants collected at baseline, 12 hours, and 7-17 days after each vaccination, as shown in Fig. 5b. After stringent quality control, a total of 515645 high-quality single-cell transcriptomes from three donors were analyzed. Clustering and annotation identified 13 major immune cell populations, encompassing lymphoid and myeloid lineages (T cells, B cells, NK cells, monocytes, and dendritic cells) (Fig. 5c and Fig. S25). Interestingly, populations of APCs, including B cells and CD14^+^ monocytes, as well as CD8^+^ T cells, exhibited a transient increase post-vaccination, returning to baseline levels within 7 days. This indicates dynamic, yet reversible, changes in cellular composition following vaccination. Strikingly, in one subject, B cells and CD14^+^ monocytes remained elevated at 30 days post-vaccination. In contrast, plasmacytoid dendritic cell (pDC) numbers gradually increased over time, but also returned to baseline by day 30. Conversely, γδ T cell (gamma delta T cell) numbers increased after the second vaccine dose and remained above baseline levels at day 30. These results implied that vaccination does, indeed, elicit an immune response in the body and that it began transitioning towards immune memory by 30 days (Fig. 5d).

To obtain a global view of gene expression changes, we quantified differentially expressed genes (DEGs) in each cell type relative to its baseline. Remarkably, hundreds of DEGs were detected in myeloid cells, T cells, and B cells within 12 hours post-vaccination (T1, T3, T5 vs. T0) (Fig. 5e), reflecting rapid and robust innate and adaptive activation. By day 7 post-first and post-second vaccination, as well as two weeks post-third vaccination (T2, T4, T6 vs. T0), transcriptional activity largely returned to baseline, indicating transient, yet repeatable, activation cycles. We next examined changes in key immune subsets. CD14⁺ monocytes displayed increased expression of HLA-DR, CXCL8, CD14, CD36, IFI30, CD68, CTSS, and TLR4, consistent with an activated, antigen-presenting and proinflammatory phenotype. Plasmacytoid dendritic cells (pDCs) also showed increased HLA-DR, FOS, CTSB, and CD74 expression, indicative of heightened interferon responses (Fig. 5f). In parallel, γδ T cells upregulated NFATC2, GRAP2, PLCG1, IL2RB, and CARD11, suggesting greater TCR signaling, cytokine responsiveness, and proliferative potential. CD4⁺ T cells upregulated JUNB, CFL2, FOS, and AKT3, reflecting robust effector activation supported by metabolic reprogramming and cytoskeletal remodeling (Fig. 5g). These signatures demonstrate that Apt-circRNA-KR2 promotes functional activation of both unique γδ T cells and conventional CD4⁺ T cells, reinforcing coordinated cellular immunity against KRAS antigens.

To resolve pathway-level dynamics, we performed gene set enrichment analysis (GSEA) using Kyoto Encyclopedia of Genes and Genomes (KEGG) pathways across major immune subsets (Fig. 5h). This analysis revealed broad induction of innate and adaptive signaling programs, most prominently at T1, T3, and T5, coinciding with peaks in DEG responses. Pathways related to antigen processing and presentation, TCR/BCR signaling, Th1/Th17 differentiation, and NK cell-mediated cytotoxicity, together with TLR, NF-κB, JAK-STAT, and RIG-I-like receptor pathways, were highly enriched across lymphoid and myeloid lineages, highlighting coordinated activation of both adaptive and innate immunity. Proinflammatory modules, including TNF, chemokine, and IL-17 signaling, were also upregulated, supporting the establishment of an inflammatory milieu favorable for T-and B-cell priming. Importantly, interferon-related pathways remained activated on day 7, suggesting a sustained antiviral-like response that may provide a favorable therapeutic window for enhancing KRAS-targeted immunity.

Finally, to assess whether vaccine-induced interferon responses target KRAS mutations, PBMCs collected at baseline and day 7 post-vaccination were stimulated ex vivo with G12D and G12V peptides (Fig. 5i). At baseline and day 7 post-first vaccination, antigen-specific IFN-γ responses were undetectable. Strikingly, however, on day 7 after the second vaccination, we saw that the vaccine elicited the highest frequencies of IFN-γ-secreting cells in response to both peptides, consistent with the enhanced activation of pDCs and CD4⁺ T cells. In contrast, on day 7 post-third vaccination, we observed reduced peptide-specific IFN-γ responses, suggesting that overactivation of pre-existing immunity may dampen subsequent boosting. By day 30, IFN-γ responses had diminished, confirming that Apt-circRNA-KR2 induces controlled, but transient, antigen-specific T cell responses.

## Discussion

Large-scale clinical studies have confirmed that the LNP delivery system in mRNA COVID-19 vaccines exhibit off-target effects, which may lead to primary or secondary myocarditis/pericarditis in some vaccine recipients^(52)^. Consequently, addressing the off-target effects of mRNA delivery systems has occupied the focus of researchers across disciplines. Recent studies concentrate on optimizing carrier formulations or incorporating targeting ligands to achieve targeted delivery of functional mRNA to specific organs or cells of the host. However, most of these approaches typically require the introduction of additional components. This not only increases the complexity of the manufacturing process, hindering large-scale production, but it also involves the introduction of extra moieties that may raise even more complex safety concerns. Addressing these issues may advance the development of mRNA-based therapeutics, especially for the therapy of pre-existing diseases, such as cancer.

Here, we report a carrier-free, scalable, and programmable aptamer-embedded circular RNAs (Apt-circRNAs) design which confers targeting capability through the strategic incorporation of diverse aptamers at varying densities, while retaining circular RNA stability and protein production capability. By embedding two distinct aptamers i.e., min2, which directs the construct to DC-SIGN on dendritic cells, and waz, which promotes cytoplasmic uptake *via* transferrin receptor, within the circRNA framework, we achieved efficient lymph node accumulation and sustained antigen expression with minimal off-target organ exposure. The self-adjuvanting nature of Apt-circRNA vaccines activates multiple innate immune-sensing pathways, including TLR1/2, TLR7/8, and RIG-I, leading to robust inflammatory signaling and antigen presentation comparable to that of LNP-delivered circRNA. Notably, the vaccine induced potent CD8⁺ and CD4⁺ T cell responses against a range of antigen types and demonstrated significant antitumor activity across five immunocompetent murine models, primarily mediated by cytotoxic T cells. Beyond directly activating antigen-specific T cells, Apt-circRNA also remodels the tumor microenvironment (TME) by inducing autophagy and apoptosis in cancer-associated fibroblasts, underscoring its multimodal mechanism of action. Promisingly, the vaccine also showed efficacy in a humanized mouse model of colorectal cancer, supporting its potential translatability, despite the inherent limitations of such accelerated tumor models.

In a first-in-human trial, the Apt-circRNA vaccine demonstrated a highly favorable safety profile. Among participants in dose-escalation (n=9) and repeated-administration (n=3) cohorts, the only adverse event was transient influenza-like symptoms that resolved within 12 hours in one individual. No grade ≥2 toxicities or injection-site reactions were observed, and all hematologic and immune parameters remained within normal ranges through 180 days. PBMCs scRNA-seq profiling revealed an immune response characterized by its rapid onset and coordinated nature. APCs, including B cells, CD14⁺ monocytes, and CD8⁺ T cells, peaked at 12 hours post-vaccination and returned to baseline within 7 days. Notably, one subject exhibited a sustained elevation of B cells and monocytes at day 30, and γδ T cell expansion was observed after the second dose. Transcriptomic profiling identified hundreds of differentially expressed genes at 12 hours, such as HLA-DR, CXCL8, NFATC2, and STAT5 across myeloid, as well as T and B cell subsets, all of which largely normalized by day 7. Gene set enrichment analysis confirmed early activation of antigen presentation, TCR/BCR signaling, and inflammatory pathways, e.g., TNF, IL-17, and JAK-STAT, with interferon response genes remaining elevated at day 7. These results demonstrate that Apt-circRNA rapidly elicits antigen-specific immunity and promotes a coordinated innate and adaptive immune response, with a favorable safety profile.

Unlike conventional platforms, Apt-circRNA requires no carrier like lipid nanoparticles or exogenous adjuvants targeting TLR or cGAS-STING pathways. Instead, subcutaneous administration directs the vaccine to target dendritic cells for transport to draining lymph nodes *via* cellular trafficking, but not as free particles, thus enabling safe and potent antitumor immunity. While this mechanism has been established in mice, further human translation will require *in vivo* imaging techniques to delineate the spatiotemporal distribution of Apt-circRNA in patients. In summary, this study establishes a solid foundation for future clinical investigation focused on aptamer-based mRNA vaccines, simultaneously providing innovative methodological insights from theoretical concepts to preclinical evaluations to accelerate the targeted delivery of other nucleic acid drugs.

### Limitations of the study

Although the Apt-circRNA vaccine platform demonstrates promising potential in our study, we acknowledge certain limitations. First, the cellular internalization efficiency of Apt-circRNA remains relatively low, which consequently affects antigen translation efficiency. To address this issue, future studies will focus on screening aptamers capable of promoting the internalization efficiency of Apt-circRNA, thereby improving antigen translation efficiency. Secondly, owing to its small size, Apt-circRNA is rapidly cleared by the kidneys in mice. This property improves safety, but it also shortens the duration of antigen translation. For tumor therapy, frequent administration would, therefore, be required. To extend the circulating half-life of Apt-circRNA, subsequent research could employ strategies that integrate other functional nucleic acid modules. Third, our study primarily used murine tumor models, which inherently differ from the human immune system, thus precluding a complete replication of immune responses in cancer patients. Furthermore, current humanized mouse models fall short in fully reconstituting myeloid cells and achieving the delivery of human-derived cancer neoantigens. Obtaining PBMCs corresponding to HLA typing for cell line-or patient-derived xenografts presents additional challenges, thus hindering the establishment of a suitable humanized immune system in mice for preclinical exploration. Last, to substantiate its antitumor efficacy and potential clinical utility, the Apt-circRNA vaccine needs to be deployed in more prospective clinical trials with a larger patient cohort to accumulate additional patient data.

## Supporting information

Methods, Fig. S1-S25, Table S1 and Table S2 will be used for the link to the file on the preprint site.

## Acknowledgments

This work was funded by the National Key Research and Development Program of China (2022YFC3401402), the National Natural Science Foundation of China (T2188102 and 22574161), and the Zhejiang Provincial Natural Science Foundation of China (LDQ23B050001 and LDQ24B020002). The authors acknowledge the support from 2024ZZBS02 of Hangzhou Institute of Medicine (HIM), Chinese Academy of Sciences. The authors acknowledge support from the Shared Instrumentation Core Facility, Hangzhou Institute of Medicine (HIM), Chinese Academy of Sciences.

## Author contributions

Y.Z., W.W. and X.Q. designed and conducted the experiments, analyzed data, and co-wrote the manuscript. J.C., X.Q., Y.D. and Z.S. designed, conducted and oversaw the human clinical trial. D.D. led the radionuclide imaging studies and analyzed data. Y.F., Z.C., Y.L. and X.Z. led and conducted sequencing and analyzed data. S.G., W.Y., H.W., K.G., Y.N., J.Z., X.K., Q.H., M.S., L.H., Y.Z., M.W., Y.Z., S.Y., C.Z., J.Z., Z.D., L.Z., T.F., Y.Z., J.J., P.Z., and X.L. conducted/assisted with experiments. S.X. and W.T. conceived and designed studies, analyzed data, provided resources, and co-wrote the manuscript.

## Data availability

The main data supporting the results in this study are available within the paper and its Supplementary Information. Any additional materials are available for research purposes from the corresponding authors upon reasonable request.

