## Supplementary material for "Programmable Aptamer-Embedded Circular RNAs for Targeted Antigen-Presenting Cells Immunotherapy": Methods, Fig. S1-S25, Table S1 and Table S2 will be used for the link to the file on the preprint site.

**The supplementary materials include:**

**Methods**

**Fig. S1-S25**

**Table S1 and Table S2**

**References**

**Methods**

**Production and purification of Apt-circRNA**

The circRNA backbone was constructed with the following components: a T7 promoter, elements from the permuted intron-exon (PIE), a spacer, aptamers, an IRES, and the coding sequence. This entire construct was synthesized by GenScript and subsequently cloned into the pUC19 plasmid. All constructs were confirmed by Sanger sequencing. Then, the plasmids were linearized by XbaI enzyme and purified by phenol/chloroform extraction. The products were concentrated to 0.5-1 μg/μl in nuclease-free water, stored at -20 °C, and used as templates for subsequent IVT. An IVT assay was carried out by using the T7 High Yield RNA Transcription Kit (NEB #E2050), following the manufacturer’s protocol. RNA was purified by LiCl precipitation after DNase I digestion for 15 min at 37 °C. To obtain circRNA *via* splicing reaction, RNA was heated at 65 °C for 3 min and then immediately placed on ice; after that, GTP (NEB) was added to a final concentration of 2 mM, and the reaction was carried out in T4 RNA Ligase Reaction Buffer (NEB #B0216L) at 55 °C for 15 min. RNA was purified by LiCl precipitation and then heated at 65°C for 3min. For every 20 μg of purified RNA, 20 U RNase R (Beyotime) and 1×RNase R reaction buffer were added to the reaction and incubated at 37°C for 2 hours. CircRNA was finally purified with the Monarch RNA Cleanup Kits (NEB #T2040 or T2050) and analyzed by 2% denaturing agarose gel electrophoresis. Especially, the Hangzhou Yunxin Zhili Biotechnology Co., Ltd. produced two distinct grades of Apt-circRNA-KR2 constructs. Research-grade materials for rat and NCG-M mouse model experiments were manufactured under strict Good Laboratory Practice (GLP) standards, and clinical-grade materials for human trials were produced following equivalent Good Manufacturing Practice (GMP) protocols.

**RNA-LNP preparation**

An ethanol phase containing all lipids and an aqueous phase of RNA were mixed to synthesize LNPs. The ethanol phase contained ionizable lipids, DSPC, DMG-PEG_2000_, and cholesterol at a molar ratio of 50: 10: 1.5: 38.5. The aqueous phase contained circRNA or mRNA in 10 mM citrate buffer. The two phases were mixed at a flow rate of 1.8 and 0.6 mL/min (3:1), respectively, using Pump33DS syringe pumps. LNPs were dialyzed in 1x PBS in a microdialysis cassette (20,000 MWCO, Thermo Fisher Scientific, Waltham, MA) at 4 ºC for 12 h. LNP diameters and polydispersity index (PDI) were measured on a Zetasizer Nano (Malvern Instruments, Malvern, UK). RNA concentration and encapsulation efficiency in LNPs were measured using a modified Quant-iT RiboGreen (Thermo Fisher) assay.

**Cell culture**

EG7.OVA cells were cultured in RPMI-1640 medium with 2 mM L-glutamine, 10% heat-inactivated FBS, 100 U/mL penicillin, 100 μg/mL streptomycin, and 0.4 mg/mL G418. DC2.4, TC-1, CT26, and HeLa cells were cultured in RPMI-1640 medium (Gibco) supplemented with 10% FBS, 100 U/mL penicillin, 100 μg/mL streptomycin, 2 mM l-glutamine, 50 μM 2-mercaptoethanol, 1× non-essential amino acids and 10 mM HEPES. B16F10, MC38, and HEK293T cells were cultured in DMEM supplemented with 10% FBS, 100 U/mL penicillin, and 100 μg/mL streptomycin. All cells were cultured at 37 °C with 5% CO_2_.

**BMDC isolation and culture**

Murine BMDCs were isolated from C57BL/6 mice (6-8 weeks). All femurs and tibiae were harvested and cut at the epiphysis level, and bone marrows were flushed out with 10 mL RPMI-1640 medium (Gibco) 3 times. The resulting cell suspensions were filtered through cell strainers (70-μm cutoff, BD Falcon), centrifuged, and then lysed with ACK buffer (Gibco). To culture BMDCs, RPMI-1640 medium supplemented with 10% FBS and 20 ng/mL GM-CSF was added to resuspended cell pellets to obtain a density of 2 x 10^6^ viable cells per 75-mm Petri dish. Three days later, an additional 10 mL of cell culture medium were added. Six days later, non-adherent and loosely adherent cells were harvested by gentle washing with PBS and then pooled for further studies.

**Cell viability assay**

BMDC cells (2 × 10^5^ cells seeded per well in a 24-well plate) were treated with Apt-circRNAs or LNP-circRNA to determine cytotoxicity. RNAs were transfected using binding buffer (DPBS supplemented with 4.5 mg/mL glucose, 5 mM MgCl_2_, 0.1 mg/mL tRNA and 1 mg/mL BSA) after seeding cells. After 6 h, the medium was replaced by complete culture medium, and the cells were incubated for another 18 h prior to adding CCK-8 reagent. After incubation for 1 h, the absorbance of cells was measured using a plate reader (Agilent BioTek) (450 nm).

**Flow cytometry**

Flow cytometry was used to analyze immunostained cultured cells, mouse blood cells, as well as tumor tissue and lymph node homogenate in single-cell suspension. Antibodies used for immunostaining were listed in **Table S1**. Cultured cells were analyzed on a BD Beckman Coulter flow cytometer (Brea, CA). Blood and tissue single cells were analyzed on the LSRFortessa X-50 (BD Biosciences). Flow cytometry results were analyzed using FlowJo V10 software.

**Antigen presentation on cultured DCs**

DC2.4 cells or BMDCs (2 × 10^5^) seeded in wells of a 24-well plate were treated with circRNA-SIINFEKL or controls for 24 h. Cells were then harvested and stained with APC-labeled anti-mouse H-2K^b^/SIINFEKL complex antibody (BioLegend). Cells were then washed and analyzed by flow cytometry.

***In vitro* cell uptake of Apt-circRNA**

*In vitro* cell uptake of dye-labeled Apt-circRNA was examined using confocal laser scanning microscopy and measured by flow cytometry. Specifically, Apt-circRNA was labeled with Cy5 *via* hybridization with a Cy5-modified 30-mer complementary DNA (cDNA). Cy5-circRNA was incubated with DC2.4 cells for a series of durations and stained with LysoTracker Green DND-26 (Life Technologies, Carlsbad, CA) and 10 μg/mL Hoechst 33342 (Life Technologies, Carlsbad, CA) for 0.5 h. Cells were then washed with DPBS three times before confocal microscopy observation on a Zeiss LSM 780 confocal microscope (Chesterfield, VA, USA). Alternatively, DC2.4 cells were seeded in wells of a 24-well plate and treated with Cy5-labeled circRNA formulations as above, followed by flow cytometric analysis as above.

**Western blot analysis**

Western blotting was performed according to standard protocols. Cell lysates were prepared using lysis buffer with protease inhibitors, and protein concentrations were determined using the BCA assay with BSA as the standard. After SDS-PAGE, proteins were transferred to PVDF membranes, blocked with 5% skim milk, and incubated with primary antibodies, as indicated in the figures. Detection was carried out with HRP-conjugated secondary antibodies, and band visualization was achieved using enhanced chemiluminescence.

**Organoid culture and transplantation**

Pathologically confirmed colorectal cancer (CRC) tissues were collected from surgical resection specimens of CRC patients. Tumor organoids were generated and maintained using the culture protocols referenced in previous studies(*1*). Briefly, tumor tissues were digested in digestion buffer (DMEM/F12 supplemented with 1.5 mg/mL Type-II collagenase, 20 μg/mL Hyaluronidase, and 10 μM Y-27632) for 30 minutes and then filtered through a 70 μm cell strainer after which the resulting single-cell suspension was collected. Following cell counting, the suspension was centrifuged, and the pellet was embedded in Matrigel (Corning, 356231) on ice. CRC organoids were cultured in complete medium (Advanced DMEM/F12 (Gibco) supplemented with 10 mM HEPES, 1× B27, 1× GlutaMAX (Gibco), 10 mM Nicotinamide, 12.5 mM N-Acetylcysteine (Tocris), 0.1 mg/mL Primocin (InvivoGen), 10 μM Y-27632 (GLPBIO), 3 μM SB202190, 10 nM Prostaglandin E2 (PGE2), 10 nM Gastrin, 0.5 μM A83-01 (Selleck), 50 ng/mL human EGF (PeproTech), 50% Wnt-3a conditioned medium (CM), 20% R-spondin CM, and 10% Noggin CM). Organoids were passaged weekly, and the culture medium was replaced every 2-3 days. For transplantation experiments, organoids were mechanically dissociated from Matrigel by pipetting, resuspended in 100 μL of complete medium containing 25% Matrigel, and slowly injected into the subcutaneous tissue of 7-15-week-old NSG mice or 10-14-week-old NCG-M mouse model.

**Animal studies**

All animal work was conducted in compliance with the Guide for the Care and Use of Animals under protocols approved by the Institutional Animal Care & Use Committee (IACUC) at the Hangzhou Institute of Medicine, Chinese Academy of Sciences. Vaccines were injected into mice subcutaneously, either into mouse footpad for vaccine delivery study or into mouse tail base for immune analysis or therapy studies. Both subcutaneous and intramuscular vaccine administrations are expected to elicit comparable magnitude and quality of antigen-specific cellular or humoral immune responses(*2*).

***In vivo* tissue and cell distribution of circRNA in mice**

To study *in vivo* tissue distribution of circRNA, it was labeled with Cy7 *via* a 30-mer cDNA. Cy7-Apt-circRNA (0.5 nmole circRNA) was *s.c.* administered in the foot pad of BALB/c mice (6-8 weeks). Days later, mice were imaged for Cy7 fluorescence using an IVIS Lumina system (Perkin Elmer). In another cohort, 6 h after administration, major organs were also isolated and imaged. The images were processed using Living Image analysis software (Perkin Elmer).

To study intranodal cell distribution of circRNA by single-cell flow cytometric analysis, BALB/c mice were *s.c.* administered at the tail base with Apt-circRNA containing fluorescently labeled circRNA (0.5 nmole Cy-5). 6 h later, draining inguinal lymph nodes were harvested, and lymph node single cells were prepared by passing minced lymph nodes through 70-μm cell strainers. Cells were then washed with and resuspended in cell staining buffer, using the following antibodies (BioLegend): Brilliant Violet 421 anti-mouse CD45, Alexa Fluor 594 anti-mouse CD11c, FITC anti-mouse CD11b, APC/Cy7 anti-mouse CD8a, PerCP/Cy5.5 anti-mouse CD4, Brilliant Violet 605 anti-mouse CD103, PE/Cy7 anti-mouse CD205, APC/Cy7 anti-mouse F4/80, PE/Cy5 anti-mouse CD11b, and APC anti-mouse NK1.1. The Zombie Aqua™ Fixable Viability Kit (BioLegend) was used to stain dead cells. Cells were then washed and analyzed by flow cytometry.

**Serum cytokine and chemokine measurement by Luminex**

C57BL/6 mice were immunized, and blood was collected under isoflurane anesthesia 12 or 24 h later. Blood was centrifuged for 5 minutes at 13,000 rpm, and sera were collected and stored at -80ºC. Preselected panels of cytokines and chemokines in the above samples were measured by Luminex (Shanghai Universal Biotech Co., Ltd).

**Tetramer staining on T cells**

Mouse CD8^+^ and CD4^+^ T cells were stained for antigen-specific tetramers using PE-conjugated tetramers (MBL International). Briefly, peripheral blood was collected from vaccinated mice, and blood cells were enriched by centrifugation. Red blood cells were lysed using ACK lysis buffer for 10 min at room temperature. Blood clots were removed using a filter. Cells were washed twice in PBS and then stained using the Zombie Aqua™ Fixable Viability Kit (BioLegend) for 20 min at room temperature. Staining was quenched, and cells were washed with FCS buffer (PBS buffer with 0.1% FBS). Cells were then blocked with αCD16/CD32 (BioLegend) for 10 min, followed by adding a staining cocktail (AF647 anti-mouse CD8a, Brilliant Violet 421 anti-mouse PD-1, and tetramer-PE) for staining at room temperature for 30 min. Cells were then washed, and 100 μL Cytofix were added into each well to resuspend cells prior to fixation at 4 °C for 20 min. Cells were then washed with Perm/Wash buffer and resuspended for flow cytometric analysis.

**Intracellular cytokine staining in T cells**

Peripheral blood was collected from immunized mice. Red blood cells were removed using ACK lysis buffer, and the obtained lymphocytes were transferred into wells of a U-bottom 96-well plate in 200 μL T cell culture media (RPMI 1640, 10% FBS, 100 U/mL pen/strep, 50 μM β-mercaptoethanol, 1x MEM non-essential amino acid solution, and 1 mM sodium pyruvate). Lymphocytes were pulsed with antigen peptides (40 μg/mL) for 4 h, followed by the addition of a BD GolgiPlug™Protein Transport Inhibitor containing brefeldin A (Thermo Fisher Scientific). The sequences of antigen peptides (GenScript) are shown in **Table S2.** Cells were then placed in a culture incubator for 6 h before incubation with αCD16/CD32 for 10 min at room temperature. Cells were stained with FITC anti-mouse CD3, AF647 anti-mouse CD8a, PerCP/Cy5.5 anti-mouse CD4 and Zombie Aqua™ Fixable Viability Kit for 20 min at room temperature. Cells were washed and subsequently fixed using Cytofix (BD Biosciences) and then washed and permeabilized in 200 μL Cytoperm solution (BD Biosciences). Next, cells were washed using Perm/Wash Buffer (BD Biosciences), and permeabilized cells were stained using PE anti-mouse IFN-γ (BioLegend) and APC/Cy7 anti-mouse TNF-α (BioLegend). Stained cells were washed for flow cytometric analysis.

**Zr-89 Radiolabeling of circRNA**

Take an appropriate amount of oxalate [^89^Zr] zirconium solution (>1 mCi) and adjust the volume to 200 μL using 1 M oxalic acid solution. Add 180 μL of 1 M sodium carbonate solution and gently mix. Then, add 1 mL of 0.5 M HEPES buffer (pH 7.1–7.3) to adjust pH to approximately 7.2. Add 40 nmol of DFO-PNA solution (2 mM, dissolved in ultrapure water), mix well, and place the reaction mixture on a constant-temperature metal heater. React at room temperature with gentle shaking in the dark for 1 h. Upon completion, purify the product using a PD-10 desalting column and determine the radiochemical purity by Radio-iTLC.

Dissolve Apt-circRNA or circRNA in PBS at a concentration of 1 mM. Mix an appropriate amount of ^89^Zr-DFO-PNA with 6 nmol of Apt-circRNA or circRNA, heat the mixture at 90 °C for 5 min, and then slowly cool to room temperature. The resulting hybridization products are ^89^Zr-DFO-PNA// Apt-circRNA and ^89^Zr-DFO-PNA// circRNA.

**PET imaging and biodistribution of Apt-circRNA.**

C57BL/6 mice (6–8 weeks) were used in PET imaging of tumor-bearing mice. Mice were anesthetized using isoflurane/O_2_ (2% v/v) before injection. Anesthetized mice were injected s.c. at the tail base with ^89^Zr-labeled vaccines (4.44−5.55 MBq/120–150 μCi each mouse) in DPBS (50 μL). At indicated time points post-injection, mice were scanned on a Trans PET Discoverist 180 (RAYCAN Technology Co., Ltd., Suzhou, China). PET images were reconstructed without correction for attenuation or scattering using the Three-Dimensional Ordered Subsets Expectation Maximization (3D-OSEM) algorithm. Inveon Research Workplace 4.2 (Siemens Medical Solutions USA, Inc.) was used for image analysis. Regions of interest (ROI) were drawn on LNs to calculate the %ID/g. The above mice were sacrificed at specified time points. Organs and blood were collected and weighed wet. The collected organs and blood, together with a series of standard solutions, were measured for ^89^Zr radioactivity on a gamma counter (Wallac Wizard 1480, PerkinElmer). The radioactivity of organs and blood was converted to calculate the percentages of the injected dose (%ID) in organs of interest and the percentages of the injected dose per gram of tissue (%ID/g).

**Tumor immunotherapy**

3 × 10^5^ EG7.OVA, MC38, TC-1, B16F10, and CT26 cells, respectively, were s.c. inoculated on the shoulder of female C57BL/6 or Balb/c mice (6–8 weeks. *n* = 6–8). For the CT26 lung metastasis model, CT26 tumor cells (3 × 10^5^) were injected intravenously into 6- to 8-week-old Balb/c mice. Tumor growth was monitored by caliper measurement. Mice were euthanized when the maximal tumor dimension reached 2 cm, the tumor volume exceeded 2000 mm^3^, or ulceration developed. Mice were treated for 1 day after tumor inoculation. Typical doses: 100 μg RNA vaccine and 100 μg αPD-1 (Bio X Cell, Inc., Lebanon, NH, USA). Vaccines were s.c. injected at the mouse tail base to allow lymphatic draining every 3 days for five times, and αPD-1 was injected *i.p.* every 3 days for five times. For lymphocyte depletion, αCD4, αCD8, and αCD20 (200 μg) were *i.p.* injected every 3 days for five times. Tumor size and mouse weight were monitored every 3 days. Tumor volumes were calculated and analyzed as described above. Results were analyzed using GraphPad Prism 7 (La Jolla, CA).

**Establishment of patient-derived organoid xenograft (PDOX) model**

To establish the PDOX model, huHSC-NCG-M mice (GemPharmatech, China) were used and housed under specific-pathogen-free (SPF) conditions in the animal facility. Mice were first irradiated, and on the following day, HSCs were administered via tail vein injection. At 7 weeks post-transplantation, mice with more than 20% huCD45^+^ cells in their peripheral blood were selected for further studies. By week 14, when both macrophages and T cells had fully differentiated, a humanized PDOX model of CRC with high KRAS G12D expression was established. One day after tumor implantation, vaccines were *s.c.* injected at the mouse tail base to allow lymphatic draining every 3 days for five times, and αPD-1 was injected *i.p.* every 3 days for five times. Typical doses: 100 μg RNA vaccine and 100 μg αPD-1 (Bio X Cell, Inc., NH). On day 20, the mice were sacrificed, and renal tumors were collected for further analysis.

**Immunohistochemistry and histological analysis**

For histological analysis, mice or rats were sacrificed after treatment with different vaccines. The heart, lung, liver, spleen, and kidneys were then collected and subjected to H&E staining, Masson’s trichrome staining, periodic acid-Schiff staining and immunostaining. These analyses were conducted to assess the morphological and pathological changes induced by the treatments.

Organoids were fixed overnight at 4°C in 4% paraformaldehyde (Biosharp) and then resuspended in molten 3% agarose, solidified at room temperature, embedded in paraffin and sectioned (4 µm). Organoids or tissue sections were deparaffinized in xylene and rehydrated through graded ethanol. For immunohistochemical (IHC) staining, antigen retrieval was performed at 95°C for 15 minutes in Tris-EDTA buffer (10 mM Tris, 1 mM EDTA, 0.05% Tween-20, pH 9.0). Endogenous peroxidase activity was blocked by incubation in 1% hydrogen peroxide solution for 15 minutes. Sections were incubated overnight at 4°C with the following primary antibodies: Ki67 (Servicebio, GB111499; 1:800), EPCAM (Servicebio, GB15274; 1:2000), and CEA (Servicebio, GB150017; 1:500). Detection used HRP-conjugated goat anti-mouse (Servicebio, G1301) IgG antibody (EPCAM, CEA) or anti-rabbit (Servicebio, G1302) IgG antibody (Ki67).

**RNA-seq**

Total RNA was isolated with the TRIzol reagent kit (Invitrogen, Carlsbad, CA, USA), following the manufacturer's protocol. The integrity and purity of RNA were assessed via an Agilent 2100 Bioanalyzer (Agilent Technologies, Palo Alto, CA, USA) with parallel verification through RNase-free agarose gel electrophoresis. Polyadenylated mRNA was enriched by Oligo(dT) magnetic beads capture. Purified mRNA underwent fragmentation and reverse transcription to cDNA using the NEBNext Ultra RNA Library Prep Kit (NEB #7530, New England Biolabs, Ipswich, MA, USA) and subsequently sequenced on an Illumina NovaSeq X Plus platform (Astrocyte Technology, Hangzhou, China). Raw sequencing reads were subjected to quality control with FastQC (v0.23.4)(*3*). Adapter removal and read quality trimming were performed on the raw reads using fastp (v0.23.4)(*4*). High-quality reads were aligned to the mouse reference genome (GRCm39) by employing the STAR aligner (v2.7.11b)(*5*). Gene abundance quantification was performed with StringTie (v2.2.3)(*6*). Differentially expressed genes (DEGs) were identified via edgeR(*7*) (FDR-adjusted P ≤ 0.05, |log₂FC| ≥ 1). Functional enrichment analyses were conducted using clusterProfiler and GSEABase R packages.

**Acute toxicity testing of Apt-circRNA-KR2 in Sprague-Dawley rats**

Seven- to eight-week-old Sprague-Dawley rats (equal number of males and females) were randomly grouped on Day 0. They received subcutaneous injections of Apt-circRNA-KR2 vaccine at different doses (250 μg, 500 μg, and 1000 μg) on Days 1, 7, and 13, respectively. All animals underwent a baseline observation prior to the first administration, followed by clinical observations and body weight measurements on Days 1, 8, 14, and 15. Blood samples were collected on Day 15 for subsequent analysis of various parameters. The entire study was conducted at Suzhou Frontage New Drug Development Co., Ltd. in full compliance with Good Laboratory Practice (GLP) guidelines.

**Long-time toxicity testing of Apt-circRNA-KR2 in Sprague-Dawley rats**

Seven- to eight-week-old Sprague-Dawley rats (equal number of males and females) were randomly grouped on Day 0. They received subcutaneous injections of Apt-circRNA-KR2 vaccine at different doses (100 μg, 250 μg, and 500 μg) on Days 1, 7, 13, and 19. All animals were observed once before the first administration (male: D-5; female: D-6). The surviving animals in the main test were subjected to detailed clinical observation once a day on D1, D7, D13, D19, D26, D33 and D40, respectively. The observation was conducted 4 to 6 hours after the administration of the drug on the administration day. Blood samples were collected on D21 and D41 for subsequent analysis of various parameters. The entire study was conducted at Suzhou Frontage New Drug Development Co., Ltd. in full compliance with Good Laboratory Practice (GLP) guidelines.

**Detection of blood routine in human blood samples**

Complete Blood Count was determined via the Mindry® BC-7800/7900 fully automated hematology analyzer (Shenzhen, China) with original Mindry® reagents (Shenzhen, China) via the electrical impedance method.

**Detection of lymphocyte subgroups in human blood samples**

Perform Peripheral Blood Lymphocyte Subset Analysis by first adding the premixed antibody cocktail (**Table S1,** volume as specified) to each tube, followed by 50 μL of well-mixed EDTA-anticoagulated whole blood; mix and incubate protected from light at room temperature for 15 minutes. Lyse red blood cells by adding 500 μL OptiLyse C Lysing Solution (Beckman Coulter), vortex gently, and incubate protected from light for another 15 minutes. Stop lysis with 1 mL PBS, centrifuge at 1500 rpm for 3 minutes, and then discard the supernatant. Resuspend the cell pellet in 500 μL PBS and immediately acquire samples on the DxFLEX flow cytometer for analysis using CytExpert software.

**Detection of cytokines in human blood samples**

Plasma cytokine detection was performed using the Cytokine Detection Kit (Jiangxi Seager Biotechnology Co., Ltd.). Perform plasma cytokine quantification by centrifuging the bead mixture at 200 × g for 5 minutes; discard the supernatant, resuspend the beads in an equal volume of bead buffer, vortex thoroughly, and incubate protected from light for 30 minutes. Vortex the bead mixture, add 25 μL to each assay tube, and then add 25 μL of sample or serially diluted standard. Add 25 μL of fluorescent detection reagent to each tube, vortex all tubes thoroughly, and incubate protected from light at room temperature for 2.5 hours. Add 1 mL PBS, centrifuge at 200 × g for 5 minutes, discard the supernatant, and resuspend the beads in 100 μL PBS. Acquire samples on the calibrated DxFLEX flow cytometer by first performing a daily quality control (QC) run with standardization beads. Then, generate a standard curve from the standard readings to calculate cytokine concentrations in the samples.

**scRNA-seq**

CD45+ cells in tumor or spleen tissues from huHSC-NCG-M mice and human PBMCs were loaded onto a 10× Genomics Chromium X™ instrument (10x Genomics) according to the manufacturer’s recommendations. The scRNA-seq libraries were processed using the Chromium Next GEM Single Cell 5' Kit v3, and the scTCRseq libraries were made using the Chromium Single Cell Human TCR Amplification Kit (10x Genomics). Quality controls for amplified cDNA libraries and final sequencing libraries were performed using the "Qualitative DNA Kit" (Agilent). The sequencing libraries for scRNA-seq and scTCR-seq were normalized to 4nM concentration and pooled. The pooled sequencing libraries were sequenced on the Illumina Novaseq Xplus platform. The sequencing parameters were as follows: Read 1 of 28bp, Read 2 of 90bp, Index 1 of 10bp and Index 2 of 10bp.

**scRNA-seq data alignment and analysis**

We demultiplexed and barcoded the sample based on the hg38 reference genome by using the Cell Ranger Software Suite (v9.0.1) and a command Cell Ranger count. After obtaining each sample, the gene counts were aggregated and imported into Seurat v5.1. The quality control standard required that cells with a detected gene count of less than 300 or more than 8500 per cell, as well as a mitochondrial gene proportion of more than 15%, be excluded. Doublets were removed using DoubletFinder (v2.0.4). Normalize the expression matrix using Seurat's "NormalizeData" function (log normalization, scale factor=1e4). Use the 'FindVariable Features' method to screen 2000 highly variable genes for subsequent dimensionality reduction and clustering analysis. Perform principal component analysis (PCA) on the expression matrix of highly variable genes, select the top 30 principal components for subsequent analysis based on the ElbowPlot, and then use Harmony (Korsunsky et al., 2019) to correct batch effects. Cluster cells using the "FindNeighbors" and "FindClusters" functions (resolution typically set to 0.5). Dimensionality reduction visualization uses UMAP methods (RunUMAP) to display the distribution and clustering of cells in two-dimensional space. Screen differentially expressed genes in each cluster using the "FindAllMarkers" function and annotate cell types using known marker genes, such as CellMarker and PanglaoDB. If necessary, use automatic annotation tools, such as SingleR or scHCL, to assist in verification. For differential expression analysis and functional enrichment, we used the DESeq2 tool for intercluster or intergroup differential gene analysis. Differential genes were further enriched in GO and KEGG pathways using tools, such as clusterProfiler(*8*), to explore their biological significance. The results of analysis were visualized using such functions as Dimplot and DotPlot provided by Seurat and further refined through ggplot2 or pheatmap packages.

**Ethics approval**

This research complies with all relevant ethical regulations. All animal procedures were conducted in accordance with the Guidelines for the Care and Use of Laboratory Animals, as approved by the Ethical Committee of the Hangzhou Institute of Medicine, Chinese Academy of Sciences (approval no. AP2024-07-0005). Human samples were collected from Zhejiang Cancer Hospital, and written informed consent was obtained from all participants. The study was approved by the Ethical Committee of Zhejiang Cancer Hospital (approval nos. IRB-2024-160/IRB-2025-136).

**Clinical trials and assessments**

This trial was conducted at Zhejiang Xiaoshan Hospital, 728 Yucai North Road, Xiaoshan District, Hangzhou, 311202, China. Detailed information, including facilities and inclusion and exclusion criteria, can be found at https://www.chictr.org.cn/searchproj.html (registration number ChiCTR2500099757). This study was approved by the Zhejiang Xiaoshan Hospital Clinical Trial Ethics Committee (approval no. EC-2024120302). Clinical outcome and safety data were reported until the clinical data cut-off of March 15th, 2025. Adverse events, laboratory values, electrocardiogram (ECG), and vital signs were assessed regularly and graded according to the National Cancer Institute Common Terminology Criteria for Adverse Events, version 5.0.

**Statistical analysis**

Data represent mean ± s.e.m., unless denoted otherwise. Statistical analysis was performed in GraphPad Prism Software, version 5.0. P-values were calculated by one-way ANOVA with Bonferroni post-test, unless denoted otherwise.

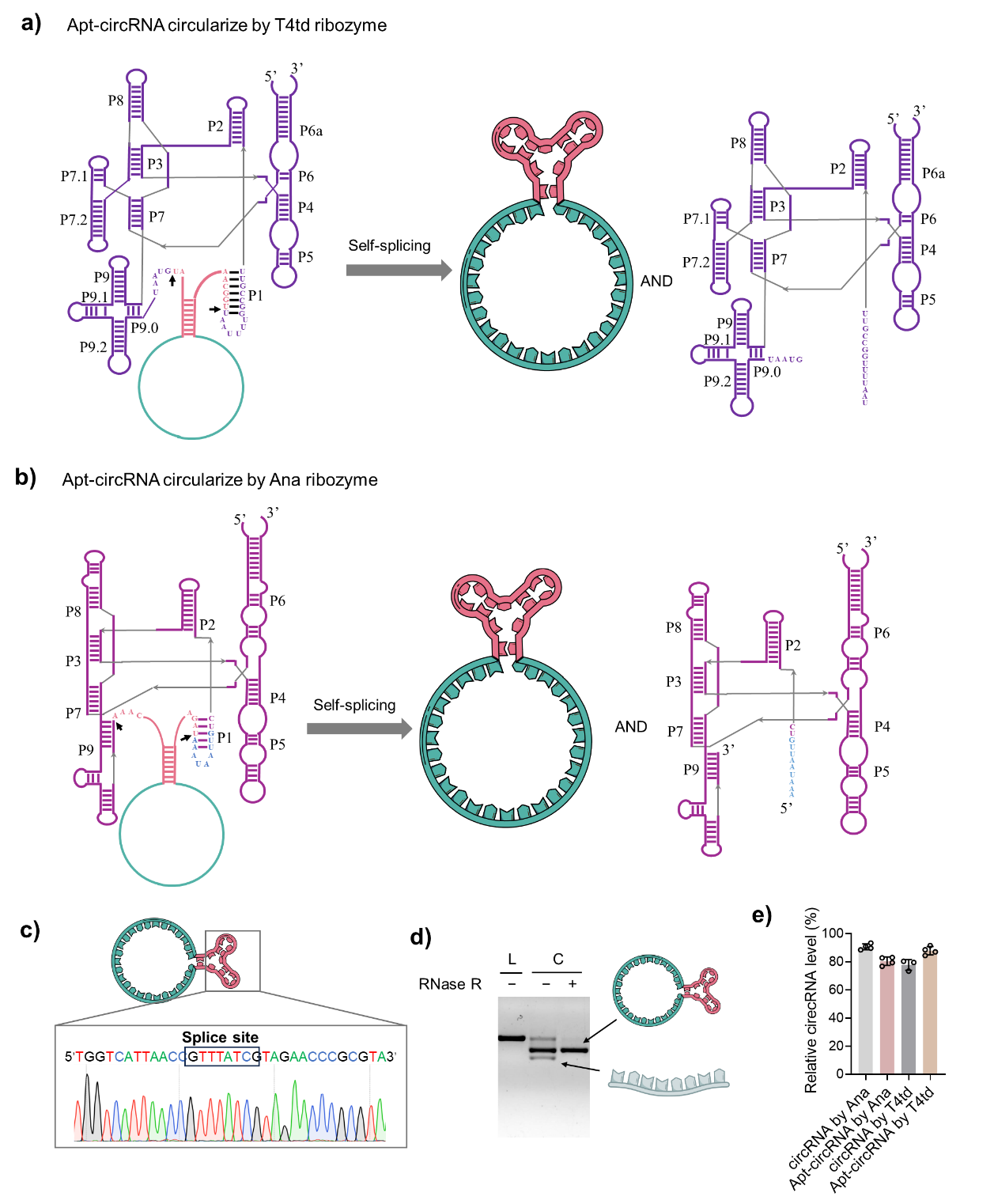

**Fig. S1. Apt-circRNAs produced by different ribozymes.**

Schematic of *in vitro* circularization of Apt-circRNAs by the phage T4 td (**a**) group I intron autocatalytic splicing and the phage Ana (**b**) group I intron autocatalytic splicing. **c**, Sanger sequencing of the cDNA of Apt-circRNA indicates precise and uniform ligation of RNA precursors into circRNA by the Ana intron-derived PIE system. The denoted RNA sequence is converted from the Sanger sequencing results of cDNA. **d,** Circularized RNA products were analyzed with denaturing agarose gel. Bands of RNA circles were verified with RNase R. **e,** Apt-circRNAs can be efficiently prepared and synthesized. Circularized RNA products are analyzed with a denaturing agarose gel and quantified with ImageJ. Data represent mean ± s.e.m., n=4.

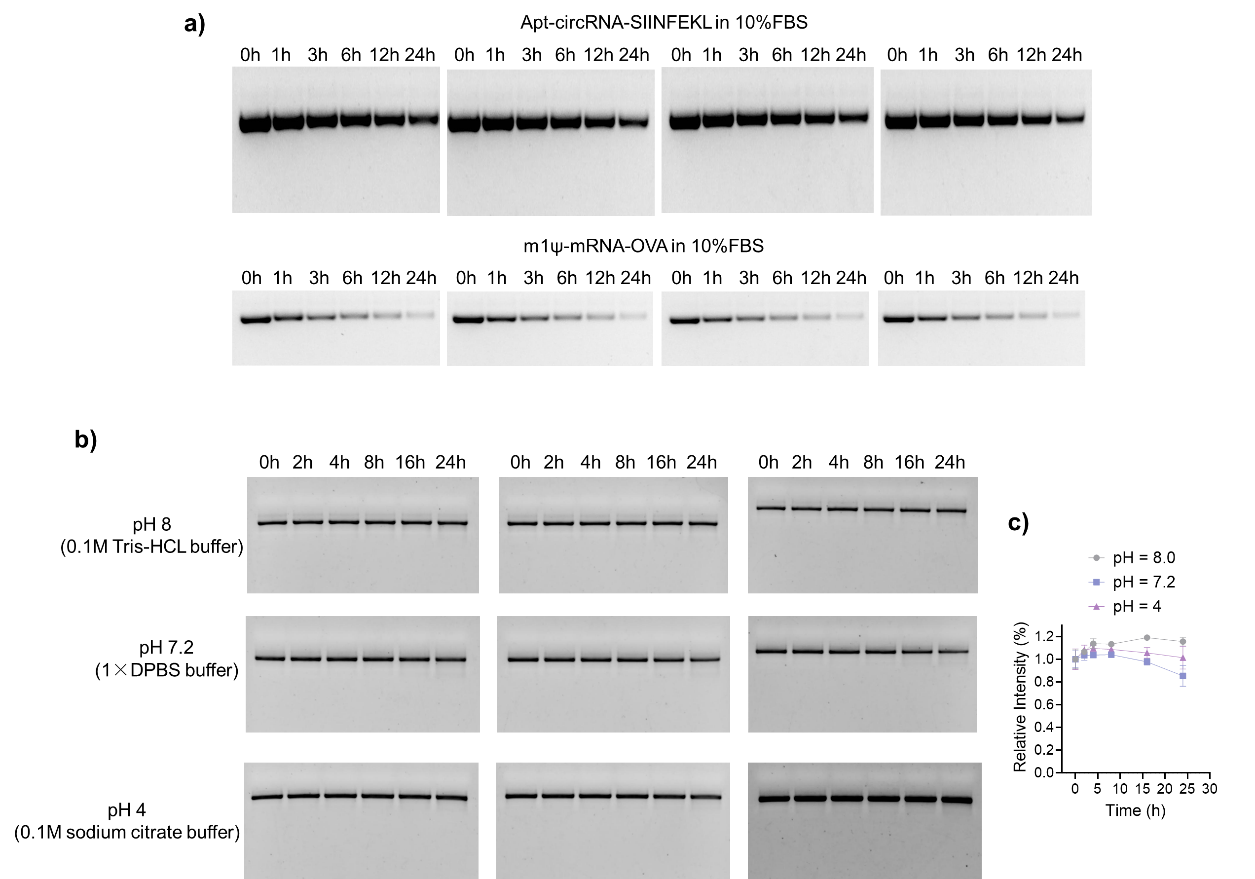

**Fig. S2. Stability of Apt-circRNA in 10%FBS or in different pH.**

**a,** The gel electrophoresis of Apt-circRNA-SIINFEKL and m1Ψ-mRNA-OVA (0.3 mg/mL) after incubation for different times in 10% FBS in DPBS (37 °C). **b,** The gel electrophoresis of Apt-circRNA (0.3 mg/mL) after incubation for different times in 0.1M Tris-HCL buffer (pH = 8.0), 1×DPBS buffer (pH = 7.2) and 0.1M sodium citrate buffer (pH = 4.0) (37 °C). **c,** Intact circRNA percentages of Apt-circRNA-SIINFEKL (0.3 mg/mL) after incubation for different times in 0.1M Tris-HCl buffer (pH = 8.0), 1×DPBS buffer (pH = 7.2) and 0.1M sodium citrate buffer (pH = 4.0) (37 °C). Data were quantified from gel electrophoresis by normalizing the band densities of different times to that of 0 hour (t-test). Data represent mean ± s.e.m., n=3.

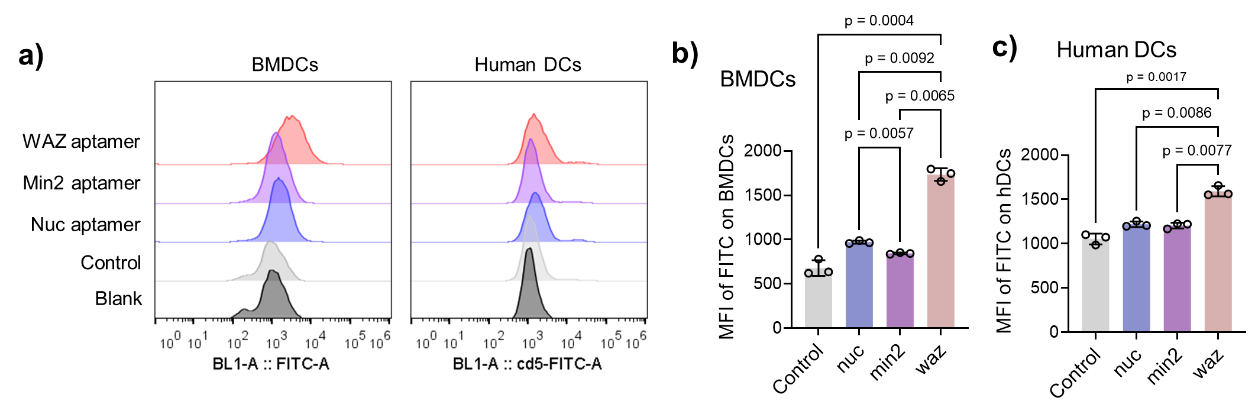

**Fig. S3. The binding of aptamers to DCs.**

**a,** Flow cytometry results of BMDCs and human DC cells incubated with different aptamers at 4 °C for 30 min. **b-c,** MFI of the aptamer in BMDCs (**b**) and human DCs (**c**). Data represent mean ± s.e.m., n=3.

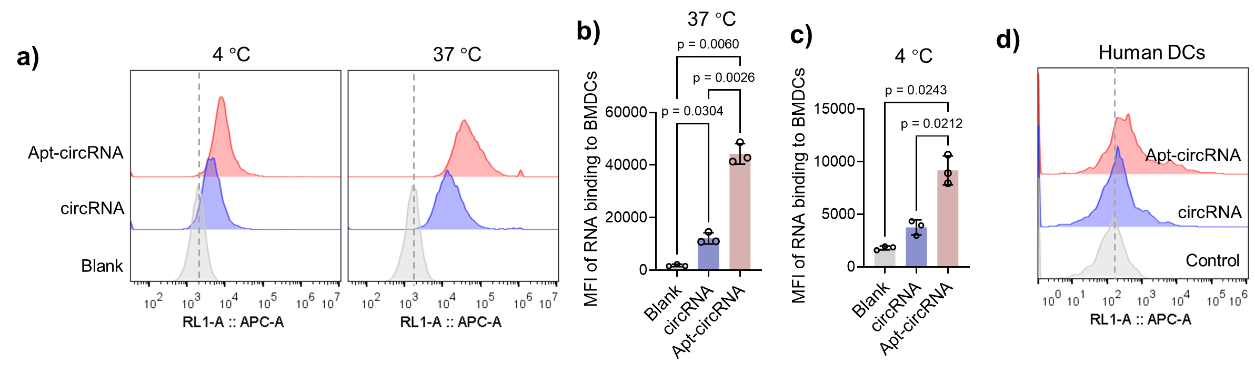

**Fig. S4.** **The binding performance of Apt-circRNA to DCs.**

**a,** Flow cytometry results of BMDCs incubated with Apt-circRNA and circRNA at 4 °C or 37 °C for 30 min. **b,** MFI of Apt-circRNA at the surface of BMDC at 37 °C. **c,** MFI of Apt-circRNA at the surface of BMDC at 4 °C. Data represent mean ± S.D., n=3. **d,** Flow cytometry results of human DCs incubated with Apt-circRNA and circRNA at 37 °C for 30 min.

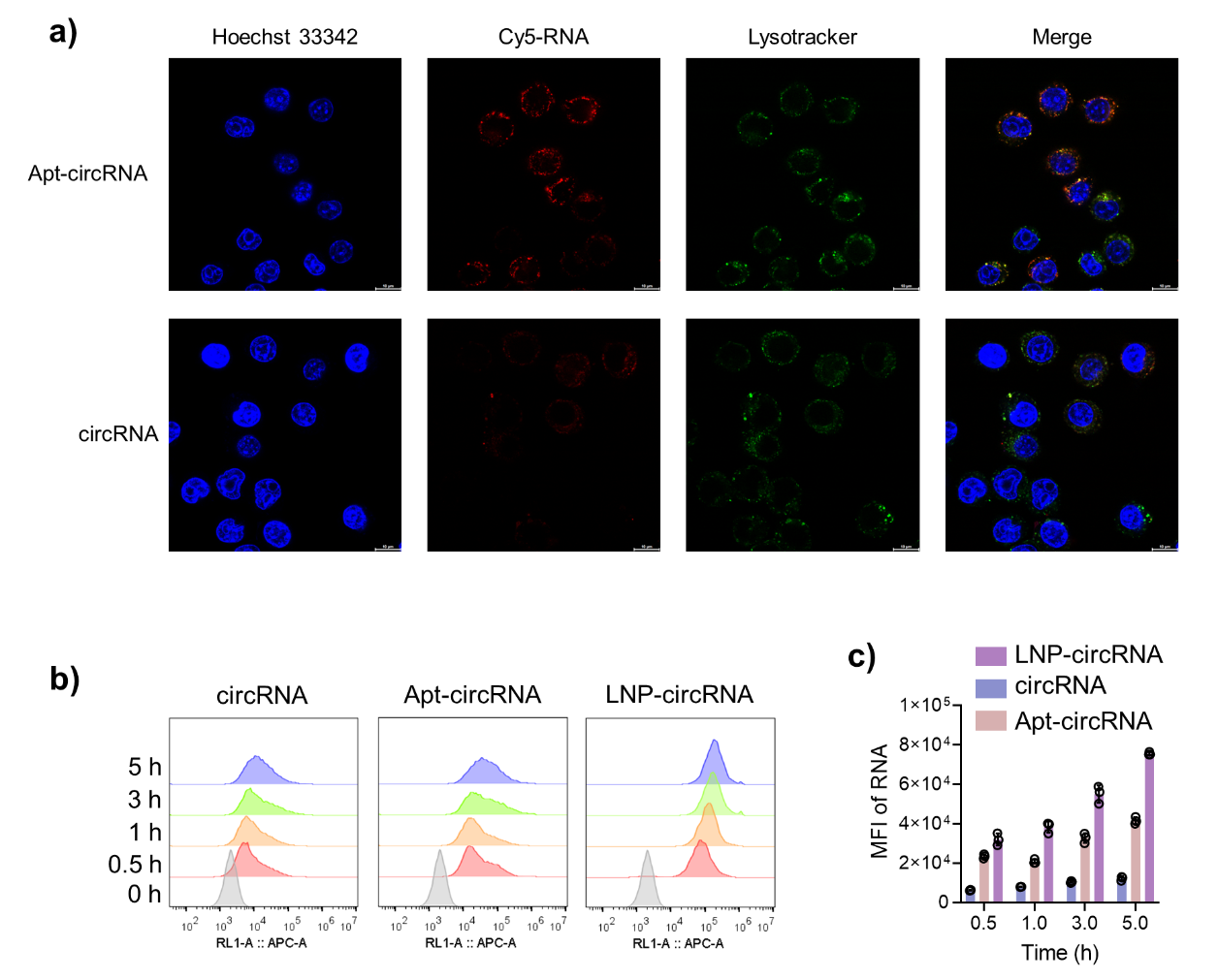

**Fig. S5. In vitro intracellular delivery of Apt-circRNA in DCs.**

**a,** Confocal microscopy images of DC2.4 cells treated with Apt-circRNA or circRNA for 4 h. Blue: nuclei stained with Hoechst 33342. Green: endolysosome stained with LysoTracker Green DND-26. Red: Cy5-circRNA. **b,** Flow cytometry results of DC2.4 cells incubated with Apt-circRNA and circRNA for different times. **c,** Flow cytometry results showing MFI of DCs incubated with Apt-circRNA or circRNA. Data represent mean ± S.D., n=3.

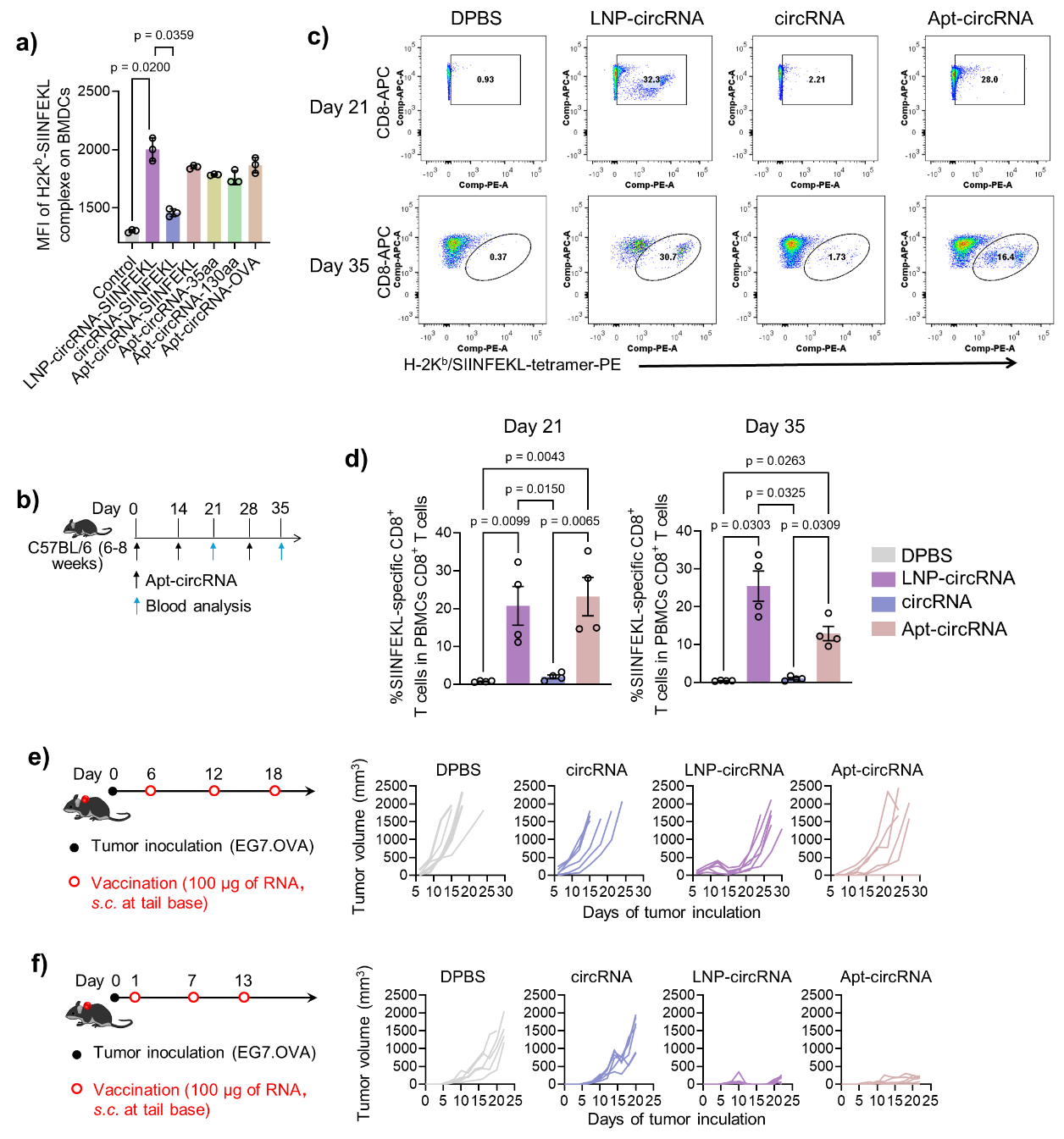

**Fig. S6.** **Apt-circRNA-SIINFEKL elicited potent T cell responses in mice.**

**a,** Flow cytometric quantification of the mean fluorescence intensity (MFI) of SIINFEKL/H-2K^b^ complexes on BMDC cells treated with Apt-circRNA coding for antigen of different lengths, LNP-circRNA-SIINFEKL, and controls, respectively, for 24 h. Data represent mean ± S.D., n=3. **b,** Design of T cell response of Apt-circRNA vaccines in vivo, using SIINFEKL as a model antigen and SM102 LNP-circRNA as a control. RNA: 100 μg, s.c. injection at the tail base of C57BL/6 mice (n = 5) on day 0, day 14, and day 28. **c,** Representative flow cytometry plots for the analysis of PBMC SIINFEKL^+^CD8^+^ T cells on day 21 and day 35 when stained using PE-labeled H-2K^b^-SIINFEKL tetramer. **d,** Tetramer staining on day 21 and day 35 showed that Apt-circRNA elicited a frequency of PBMC SIINFEKL^+^CD8^+^ T cells similar to that of the LNP-circRNA group. Data represent mean ± s.e.m., n=4. **e,** Left: for immunotherapy studies in mouse models of subcutaneous EG7.OVA, tumors were inoculated into the right flank of C57BL/6 mice. Vaccine: Apt-circRNA, circRNA, or LNP-circRNA (RNA = 100 μg), s.c. injection at mouse tail base on day 6, 12, and 18. Right: individual EG7.OVA tumor growth curves in C57BL/6 mice. **f,** Left: for immunotherapy studies in mouse models of subcutaneous EG7.OVA, tumors were inoculated into the right flank of C57BL/6 mice. Vaccine: Apt-circRNA, circRNA, or LNP-circRNA (RNA = 100 μg), s.c. injection at mouse tail base on day 1, 7, and 13. Right: individual EG7.OVA tumor growth curves in C57BL/6 mice.

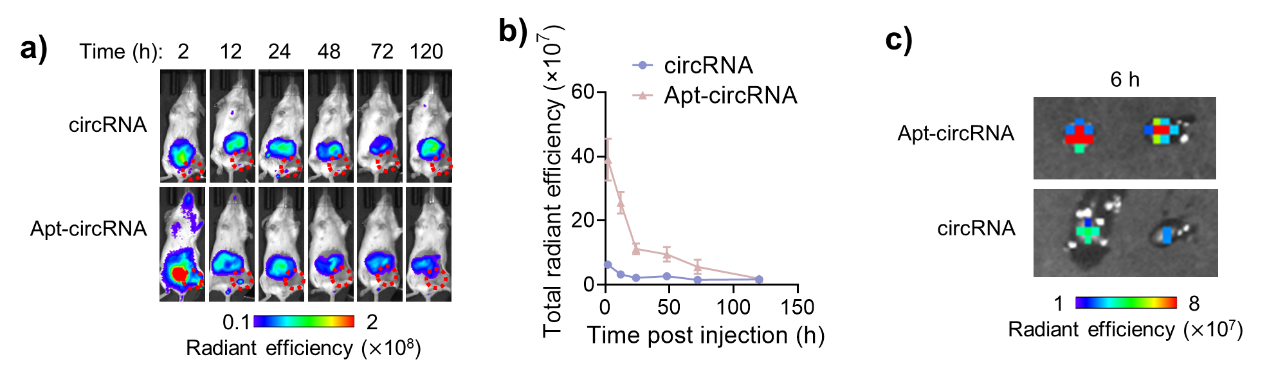

**Fig. S7. Aptamer efficiently delivered circRNA to lymph nodes.**

**a,** Aptamer promoted the delivery of Cy7-circRNA to draining popliteal lymph nodes (circled) in Balb/c mice (0.2 nmol, s.c. injection at foot pad). **b,** AUC of radiance efficiency. ***p* < 0.01 by t-test of the AUC. Data represent mean ± s.e.m., n=3. **c,** *Ex vivo* fluorescence images of draining popliteal lymph nodes after s.c. injection at tail base for 6 h.

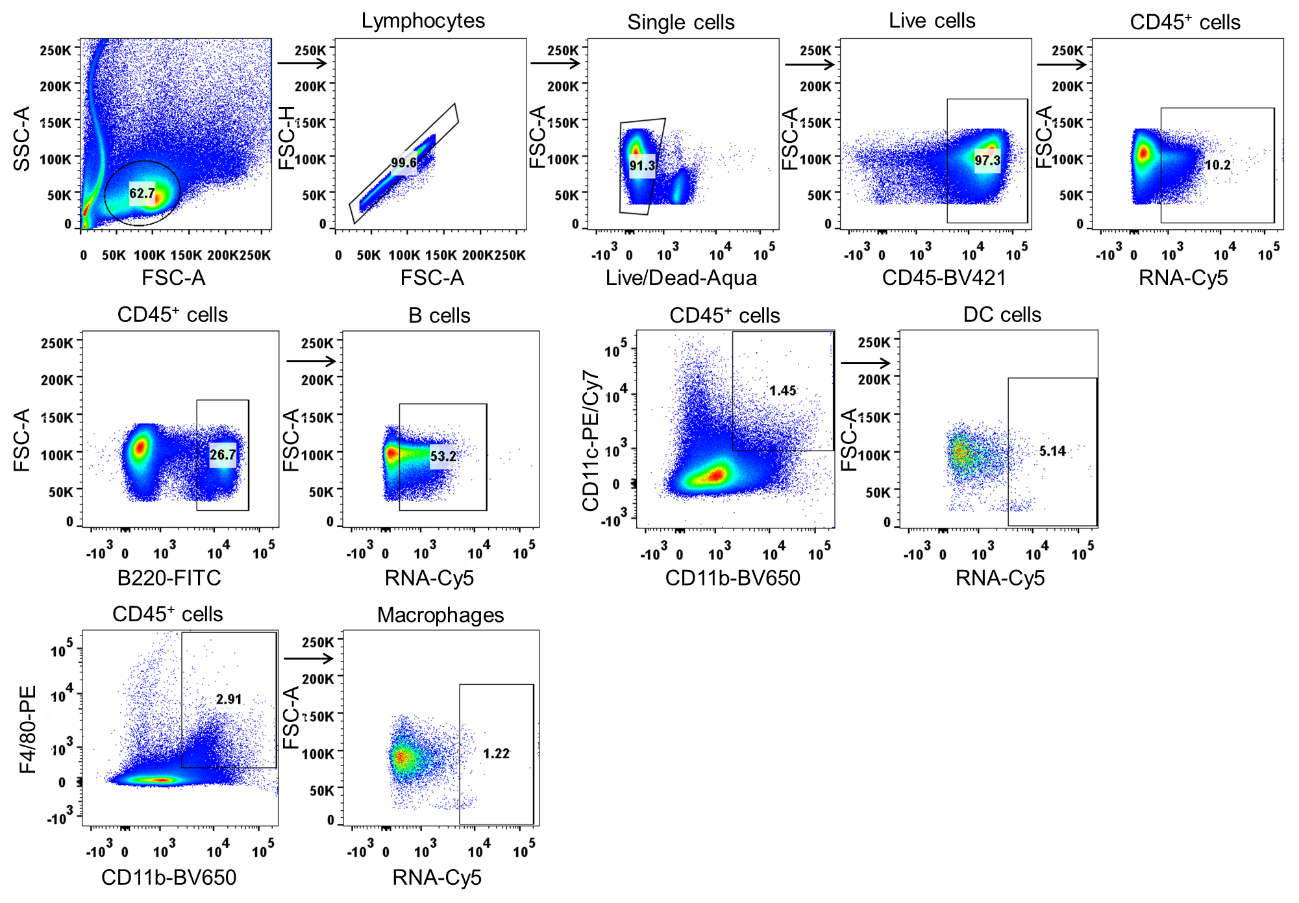

**Fig. S8.** **Representative gating tree used in flow cytometric analysis of Apt-circRNA^+^CD45^+^ cells, Apt-circRNA^+^B220^+^ B cells, Apt-circRNA^+^CD11c^+^CD11b^+^ DC cells, and Apt-circRNA^+^CD11b^+^F4/80^+^ macrophages in mouse draining lymph nodes.**

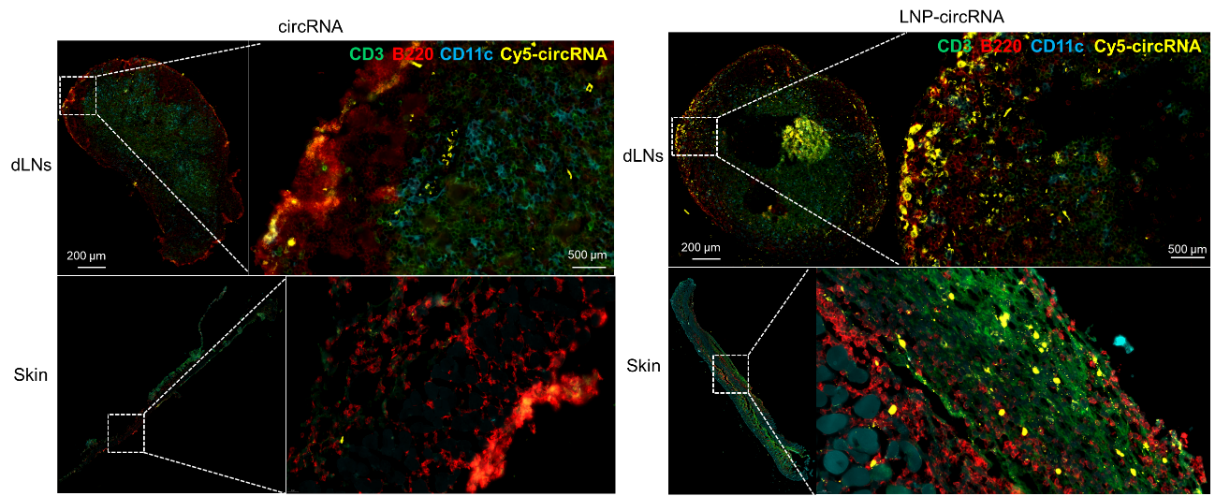

**Fig. S9.** **Immunohistochemistry of skin and inguinal LNs of mice treated with circRNA or LNP-circRNA (CD3, green; B220, red; CD11c, blue; Cy5-circRNA, yellow).**

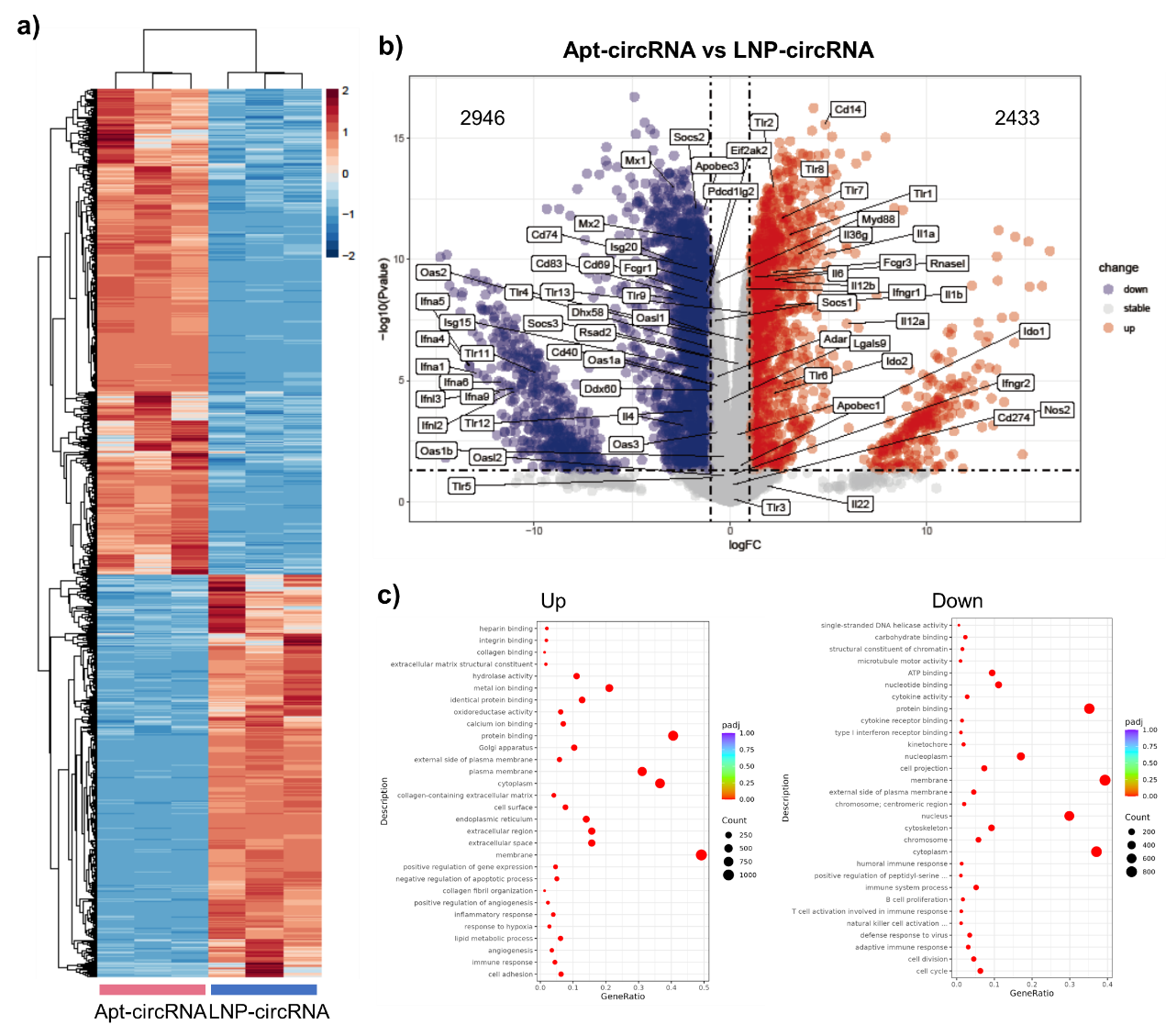

**Fig. S10. Intrinsic innate immunostimulation by Apt-circRNA.**

**a,** Transcription heatmaps of genes involved in inflammation, migration, antigen processing and presentation, TLRs, RLRs, CLRs, and miscellaneous immune-related genes. **b,** Volcano plot of differentially accessible peaks between Apt-circRNA and LNP-circRNA. **c,** Significantly enriched pathways among differentially expressed genes, as determined by GSEA.

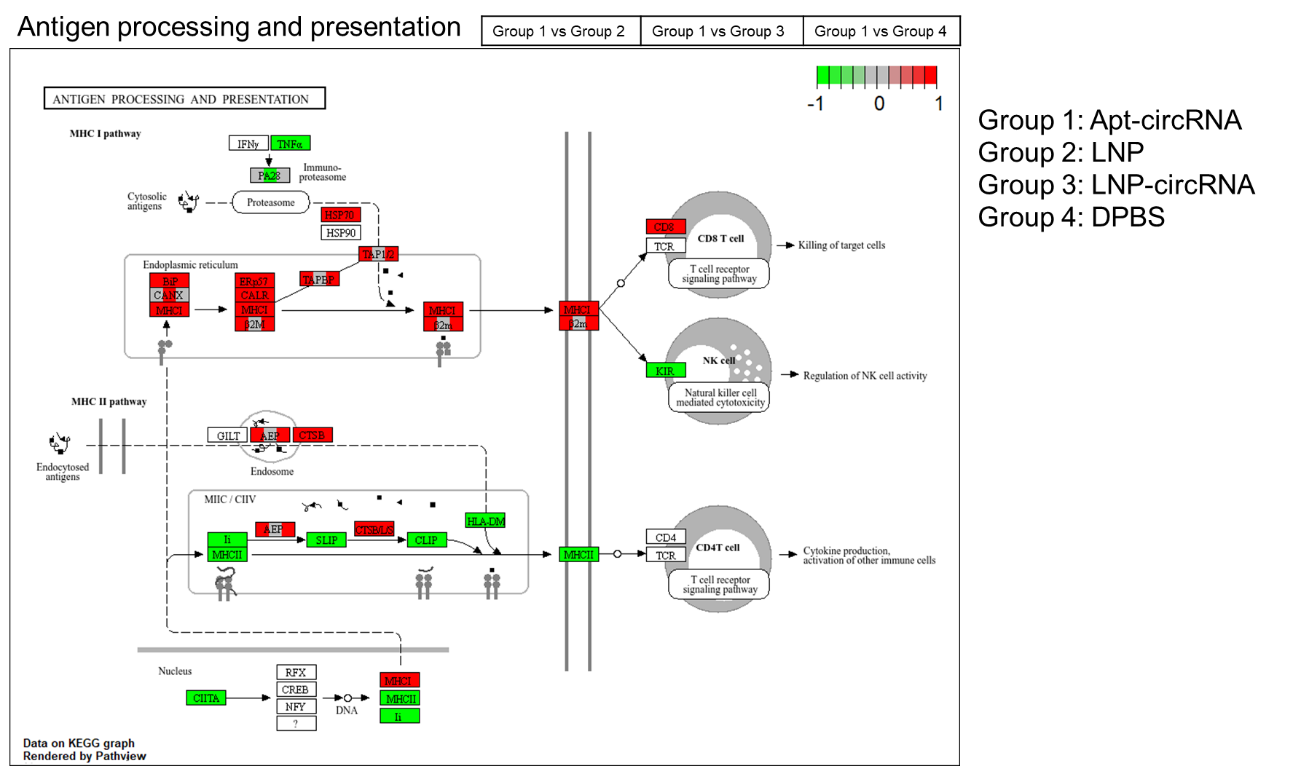

**Fig. S11. Visualization of the KEGG pathway of antigen processing and presentation by Pathview.** Gene expression values can be mapped to gradient color scale. Red indicates that Apt-circRNA induces higher gene expression levels than the other groups. Green indicates that Apt-circRNA induces lower gene expression levels than the other groups.

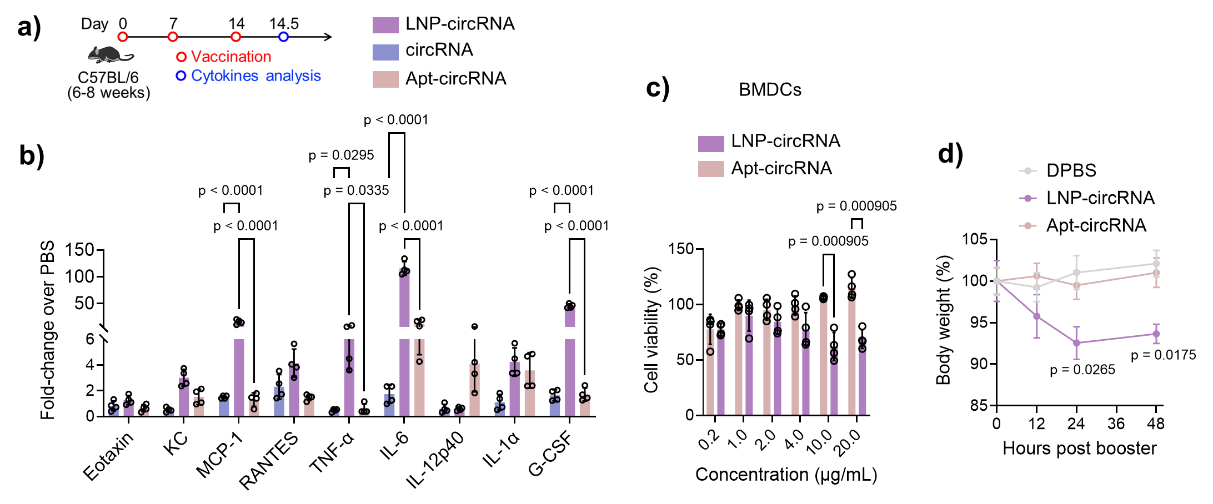

**Fig. S12. Tolerability of Apt-circRNA vaccine in mice.**

**a,** Timeline of *in vivo* safety study. C57BL/6 mice (6-8 weeks) were immunized with Apt-circRNA, as well circRNA, and LNP-circRNA, respectively. **b,** Luminex results of serum cytokines and chemokines. **c,** Cell viability of BMDCs treated with Apt-circRNA and LNP-circRNA after 24 h. **d,** Body weight of mice immunized with Apt-circRNA and LNP-circRNA at a dose of 100 μg. Data represent mean ± s.e.m., n=4.

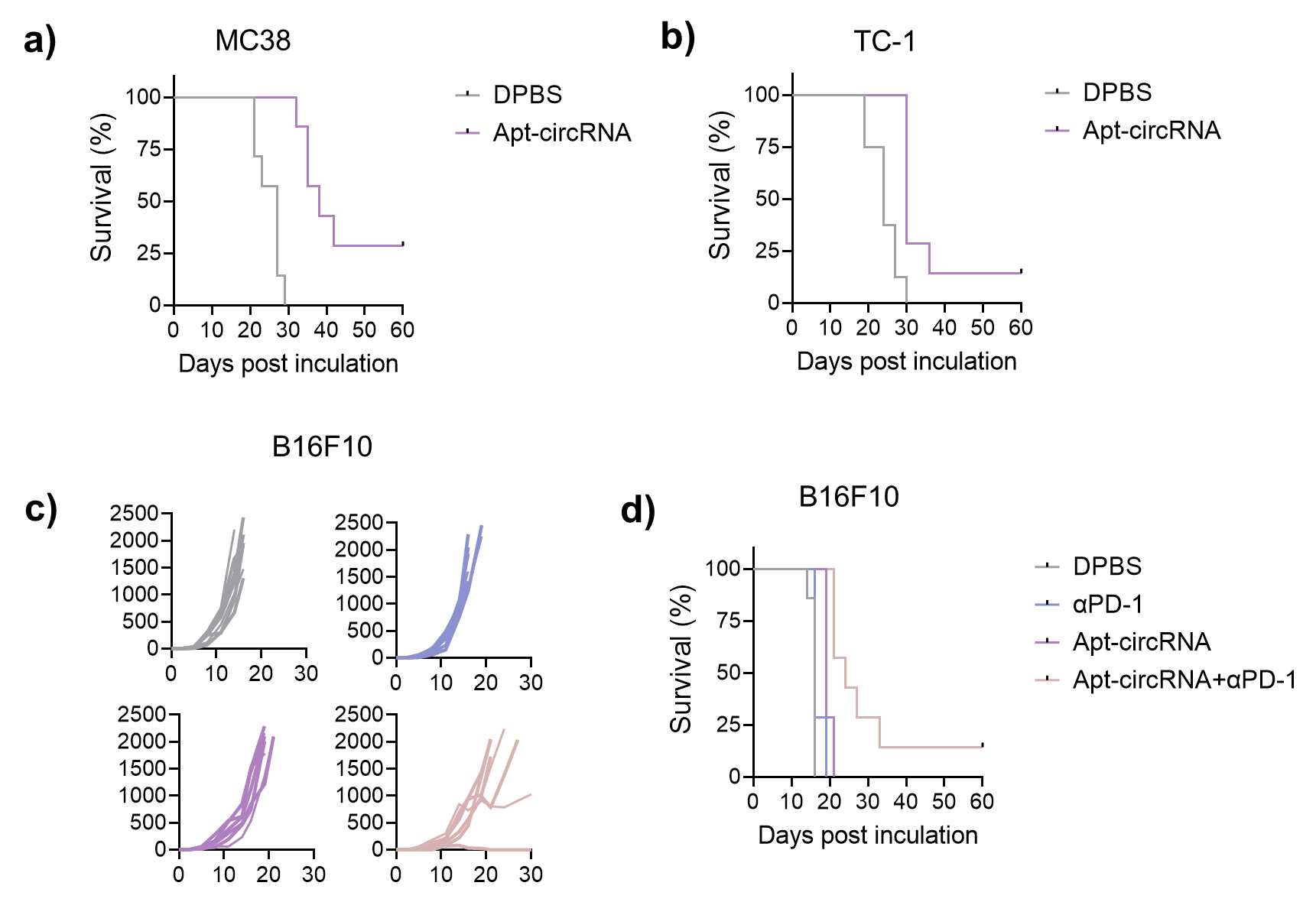

**Fig. S13. Mutivalent Apt-circRNA vaccines for robust tumor immunotherapy.**

**a-b,** Kaplan-Meier survival curves of MC38 tumor- (**a**) and TC-1 tumor (**b**)-bearing mice treated with Apt-circRNA. **c,** Individual B16F10 tumor growth curves in C57BL/6 mice treated with Apt-circRNA, as well as αPD-1 (i.p.) alone or combined with Apt-circRNA. CR: complete regression rate. **d,** Kaplan-Meier survival curves of B16F10 tumor-bearing mice treated with Apt-circRNA, as well as αPD-1 (i.p.) alone or combined with Apt-circRNA.

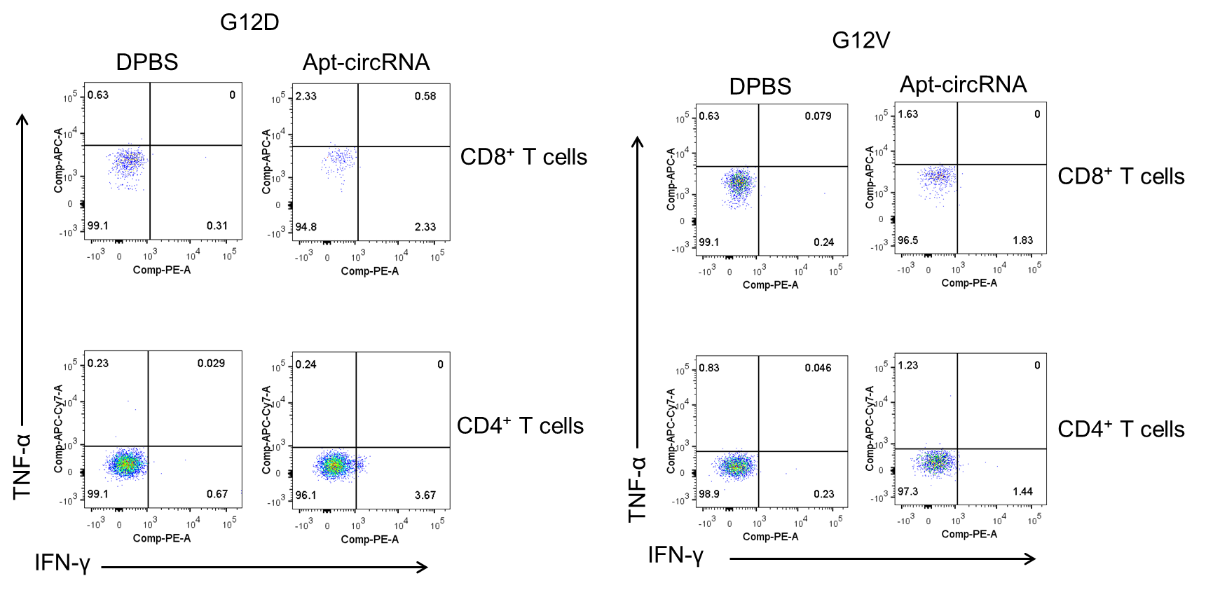

**Fig. S14. Representative flow cytometry plots of PBMC cytokines^+^CD8^+^ T cells on day 21 after three doses of Apt-circRNA (Fig. 3) when stained with PE-labeled anti-mouse IFN-γ antibody and APC/Cy7- or APC-labeled anti-mouse TNF-α antibody.**

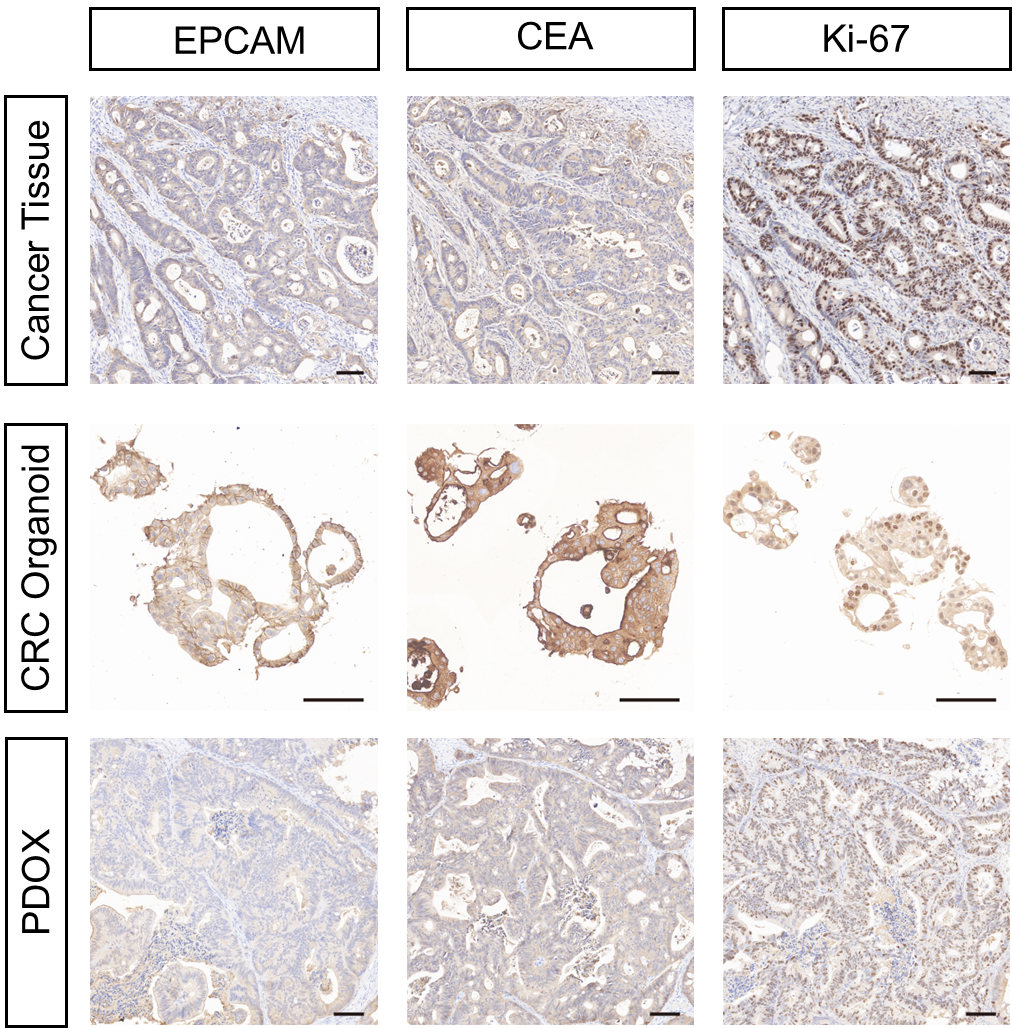

**Fig. S15. H&E and IHC staining of Patient-017 tumor tissue, organoids derived from it, and patient-derived organoid xenograft model (PDOX) derived from the organoids (PDO-017).** Organoids demonstrate morphological features similar to those of the parent tissue, including nuclear pleomorphism, acino-glandular growth pattern, and positive staining for EPCAM and CEA. They are also more proliferative than the parent tissue, as depicted by higher Ki-67 expression. Scale bar, 100 µm

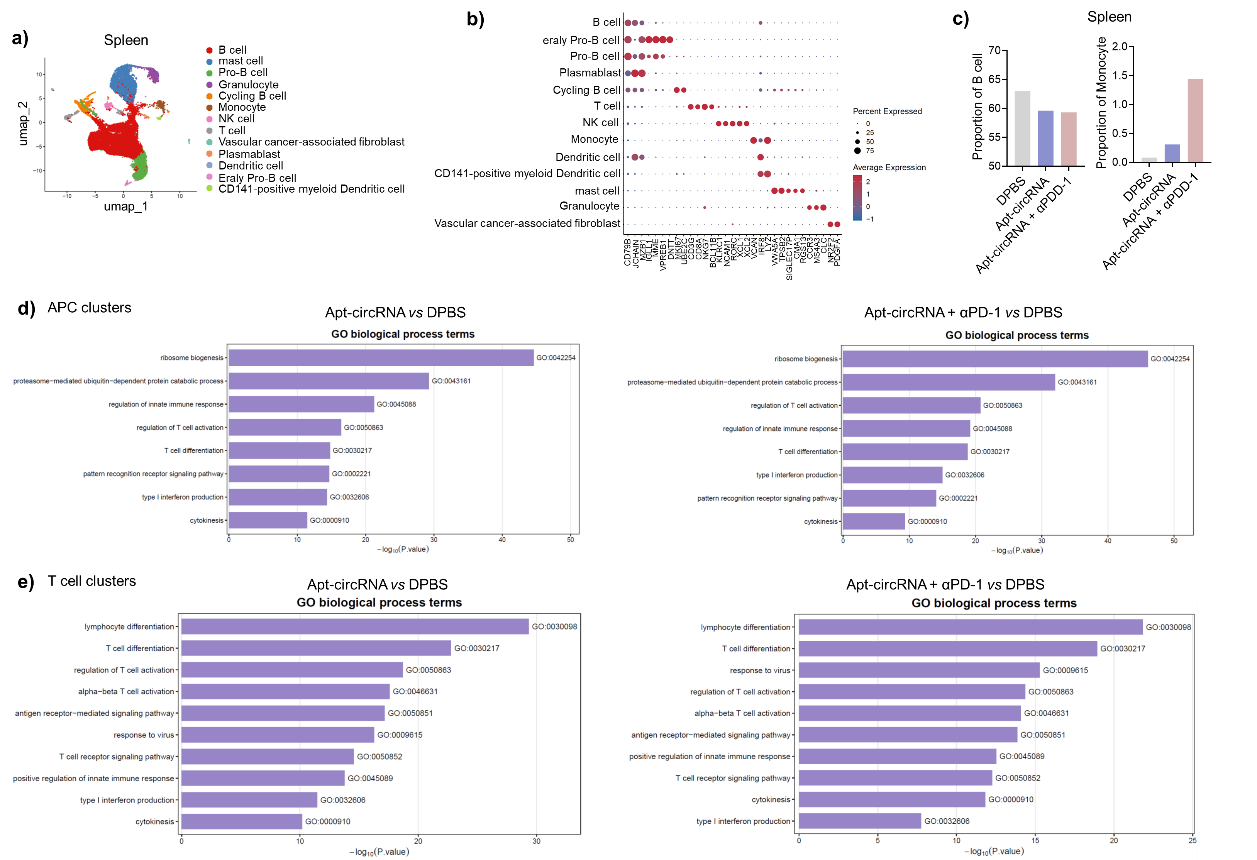

**Fig. S16. scRNA-seq data of spleen in hu-HSC-PDX CRC model.**

**a,** Uniform manifold approximation and projection (UMAP) representations of scRNA-seq data of all CD45^+^ cell subsets in spleen. **b,** Dot plot depicting cell type marker genes across 13 distinct cell types from spleen and tumor samples. **c,** Average proportion of B cells and monocytes derived from splenic tissues. **d,** Differential pathways enriched for the discriminative markers of APC cell subset in spleen by GO terms. **e,** Differential pathways enriched for the discriminative markers of T cell subset in spleen by GO terms.

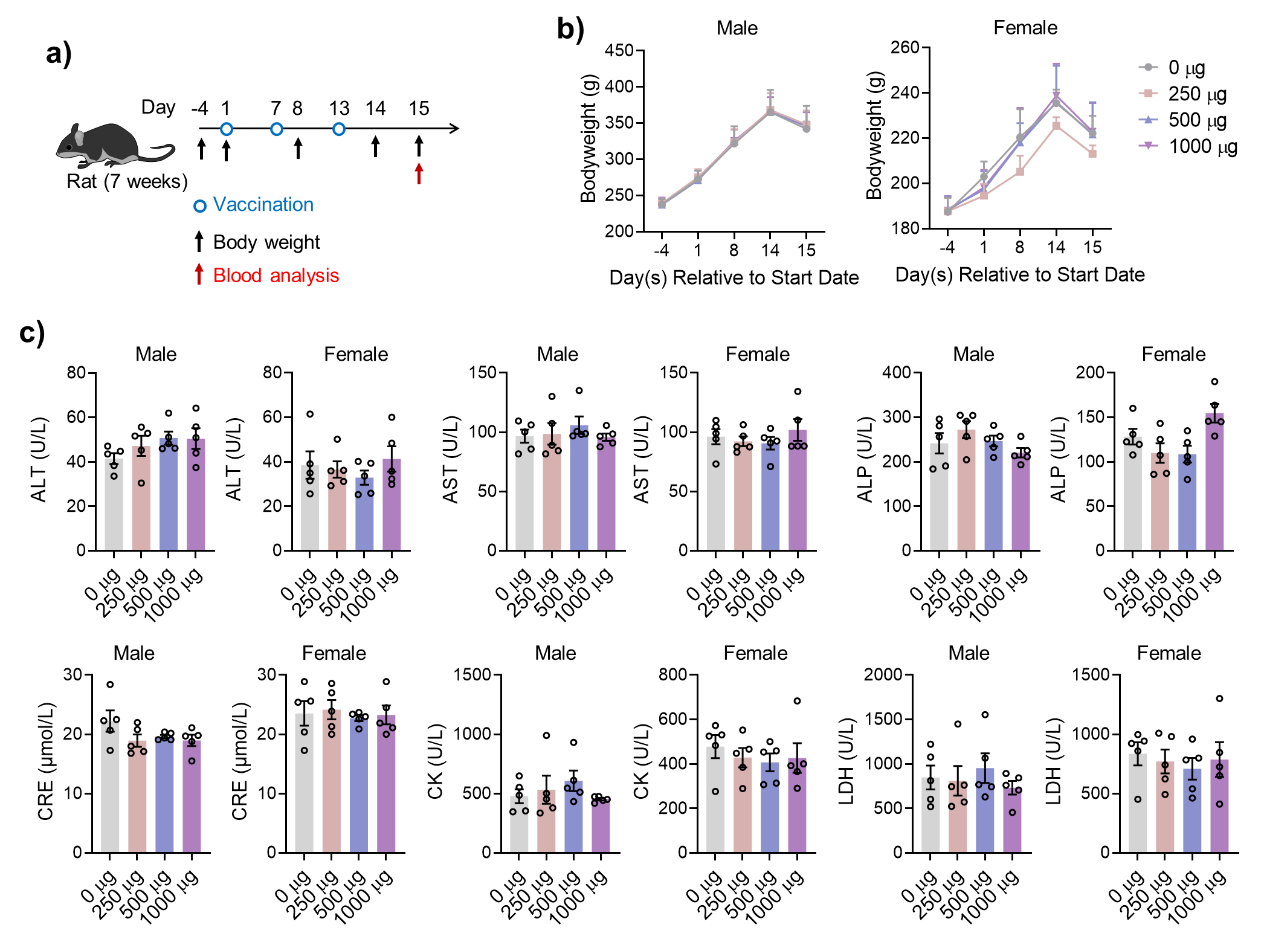

**Fig. S17. GLP-standardized acute challenge experiments of Apt-circRNA vaccine in rat.**

**a,** Timeline of *in vivo* acute toxicity study. Sprague-Dawley (SD) rats (7 weeks) were immunized with Apt-circRNA at different doses. **b,** Body weight of male and female rats before and after immunization with Apt-circRNA. **c,** Representative blood biochemical parameters in rats on day 15 in acute toxicity study. Data represent mean ± s.e.m., n = 5. *P < 0.05, **P < 0.01, ***P < 0.001, ****P < 0.0001, one-way ANOVA with Bonferroni post-test.

To systematically evaluate the *in vivo* safety profile of the Apt-circRNA vaccine in SD rats, good laboratory practice (GLP)-standardized acute challenge experiments confirmed the absence of dose-limiting toxicity for Apt-circRNA-KR2 vaccine (**Fig. S17a**). Quantitative safety evaluation revealed that all tested doses (250/500/1000 μg) of Apt-circRNA-KR2 vaccine maintained physiological stability in both male and female rats during the 14-day acute phase observation with <5% body weight fluctuation (vs. PBS control), even at the maximum dose (**Fig. S17b**). In addition, the blood biochemistry, including alanine aminotransferase (ALT), aspartate aminotransferase (AST), alkaline phosphatase (ALP), creatinine (CRE), creatine kinase (CK), and lactate dehydrogenase (LDH), did not differ between vaccinated and control rats at different doses on day 15 (**Fig. S17c**), suggesting normal liver and kidney function.

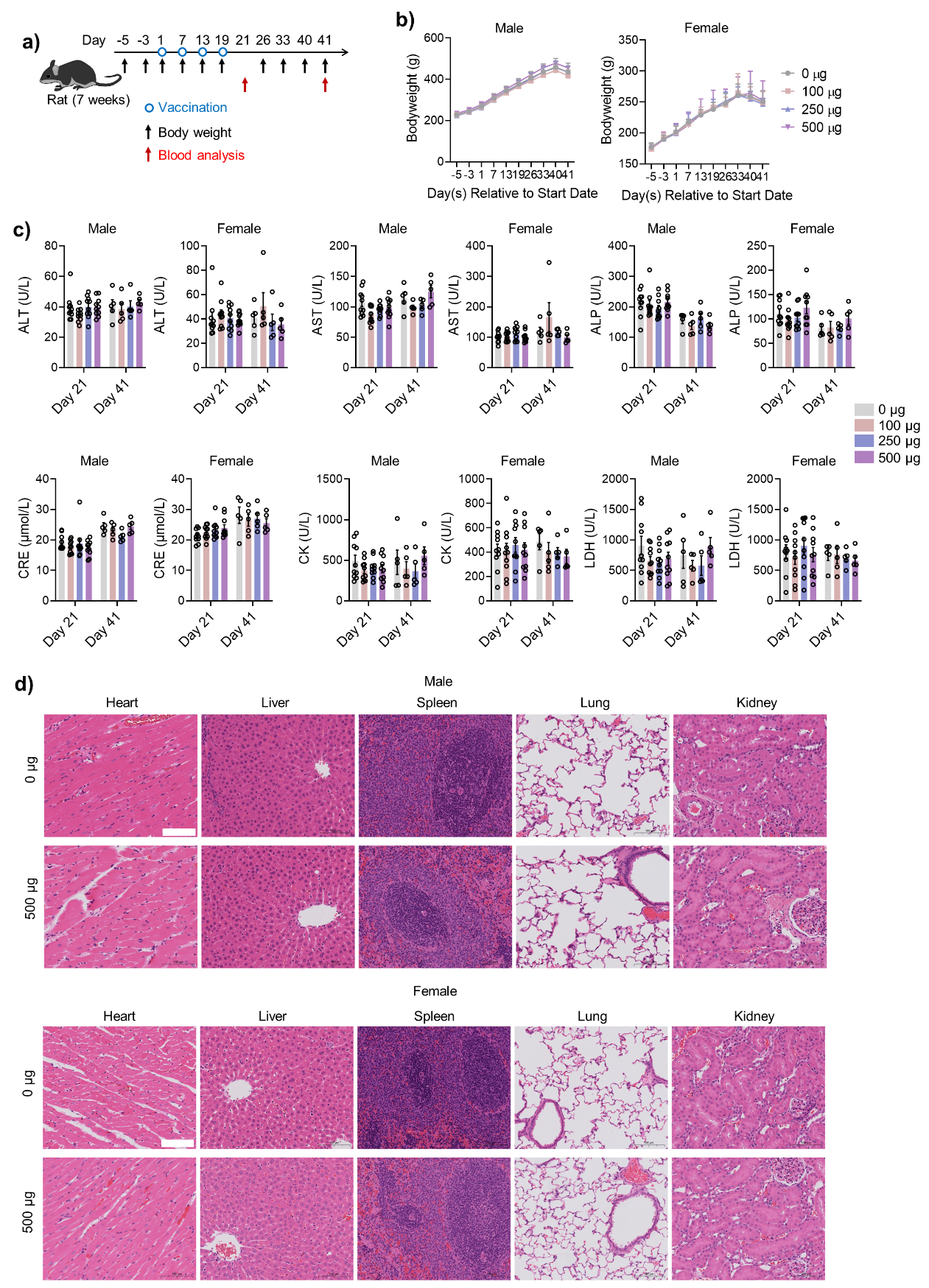

**Fig. S18. GLP-standardized systematic long-term toxicity study of Apt-circRNA vaccine in rat.**

**a,** Timeline of *in vivo* long-term toxicity study. SD rats (7 weeks) were immunized with Apt-circRNA at different doses. **b,** Body weight of male and female rats before and after immunization with Apt-circRNA. **c,** Representative blood biochemical parameters in rats on day 21 and day 41 in long-term toxicity study. **d,** Representative hematoxylin and eosin images of heart, liver, spleen, lung, and kidney of male and female rats on day 41. Scale bar, 100 μm Data represent mean ± s.e.m., n = 5. *P < 0.05, **P < 0.01, ***P < 0.001, ****P < 0.0001, one-way ANOVA with Bonferroni post-test.

We conducted a GLP-standardized systematic long-term toxicity study to test the Apt-circRNA-KR2 vaccine (**Fig. S18a**). Even in the highest dose group, results showed no significant changes in rat body weight before vaccination, after vaccination, or on days 21 and 41 post-vaccination, compared to the control group (**Fig. S18b**). In addition, analysis of blood biochemical parameters in rats on days 21 and 41 post-vaccination revealed no difference between any vaccinated group at any dose and the control group, indicating normal liver and kidney function (**Fig. S18c**). Last, hematoxylin and eosin staining of major organs on day 41 did not reveal discernible pathologies or inflammatory lesions in rats that received four doses of Apt-circRNA vaccine (**Fig. S18d**). Taken together, these findings demonstrate a satisfactory safety profile of this Apt-circRNA vaccine in rat models.

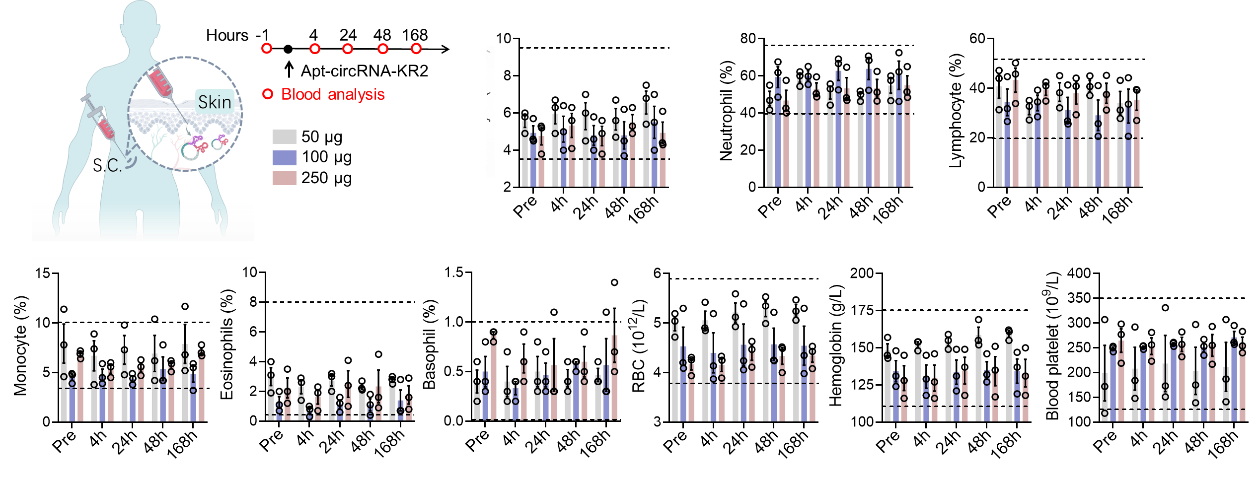

**Fig. S19. Tolerability of Apt-circRNA vaccine in human.** The hematologic parameters of 9 healthy volunteers after immunization with a single dose of Apt-circRNA-KR2 vaccine (50, 100, and 250 μg; n=3/group). Black dashed line represents the range of normal individuals. Data represent mean ± s.e.m.

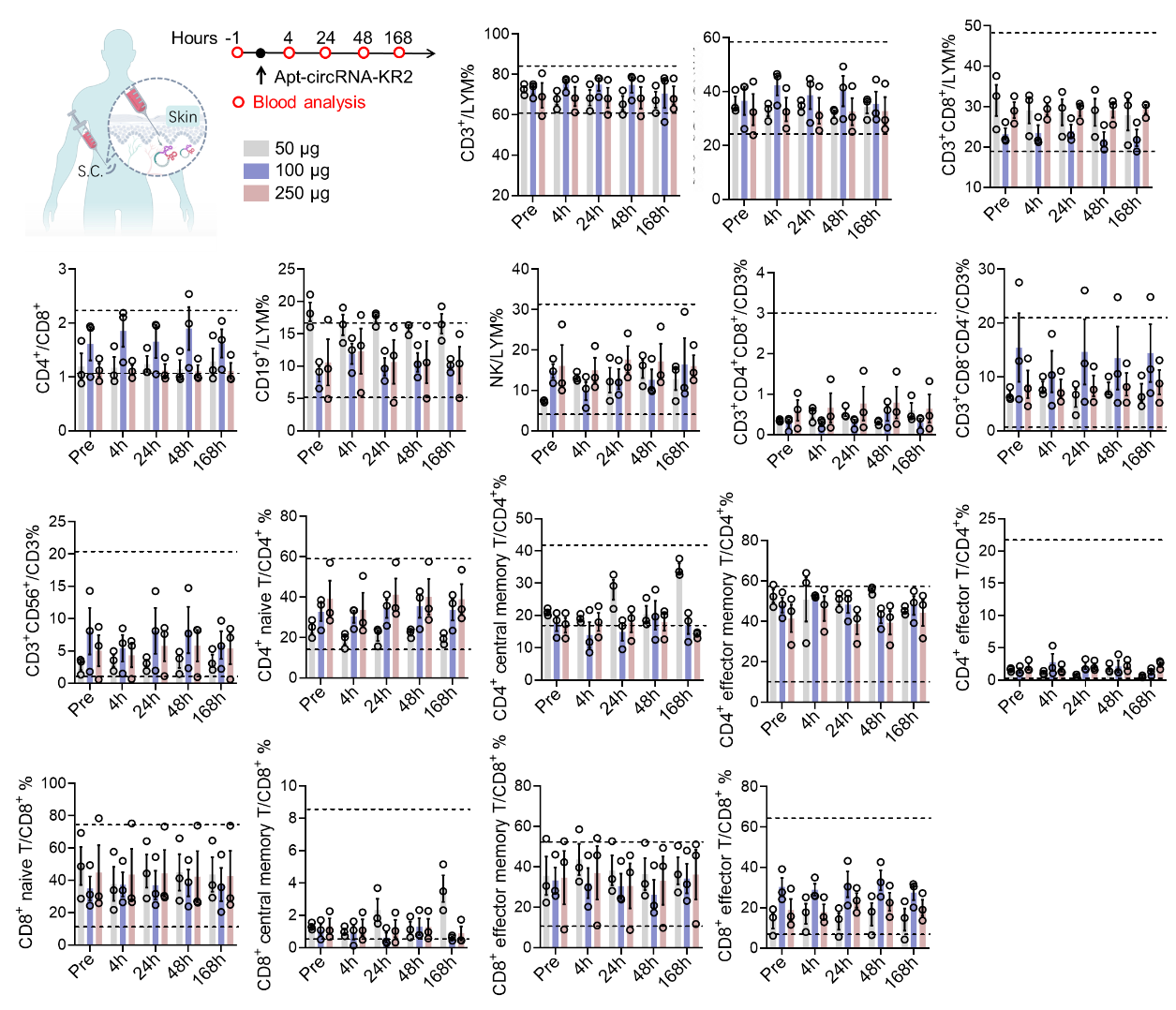

**Fig. S20. Tolerability of Apt-circRNA vaccine in human.** The immune cell subsets of 9 healthy volunteers after immunization with a single dose of Apt-circRNA-KR2 vaccine (50, 100, and 250 μg; n=3/group). Black dashed line represents the range of normal individuals. Data represent mean ± s.e.m.

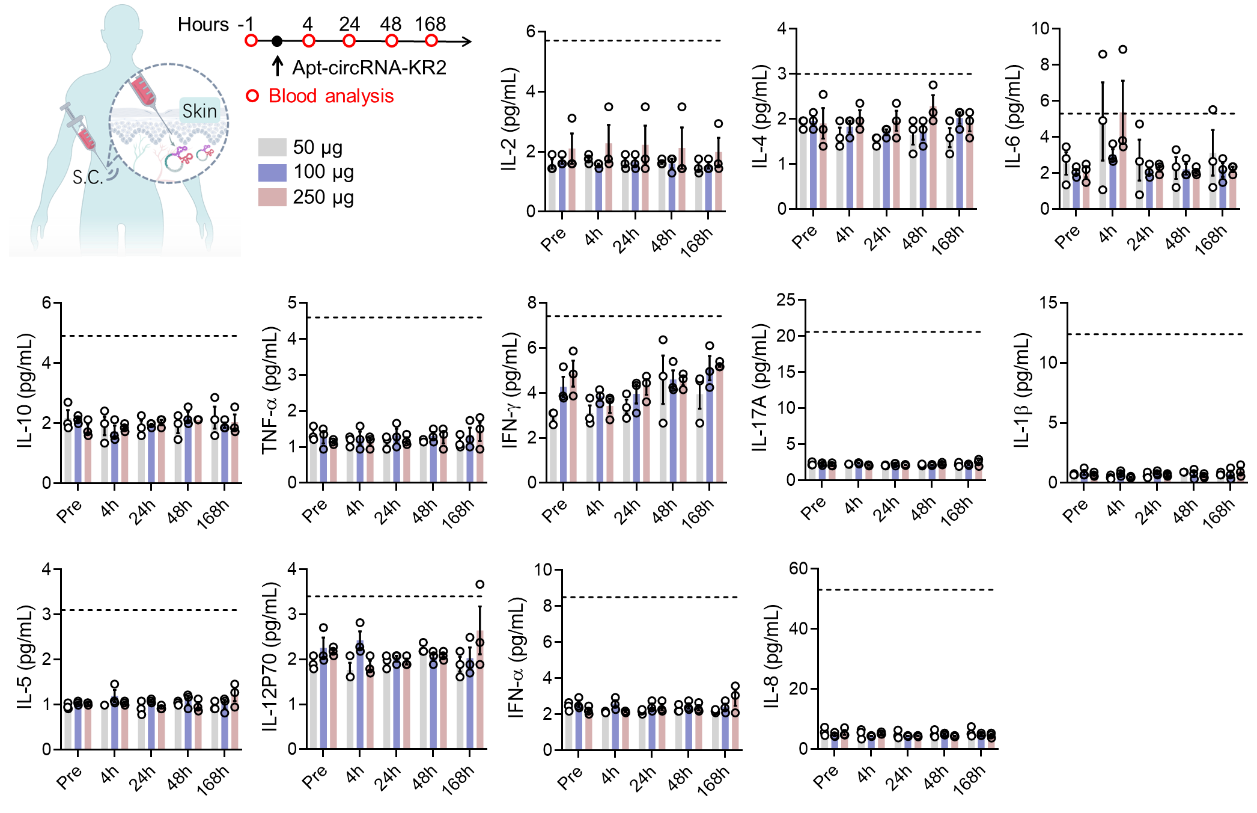

**Fig. S21. Tolerability of Apt-circRNA vaccine in human.** The serum cytokines of 9 healthy volunteers after immunization with a single dose of Apt-circRNA-KR2 vaccine (50, 100, and 250 μg; n=3/group). Black dashed line represents the range of normal individuals. Data represent mean ± s.e.m.

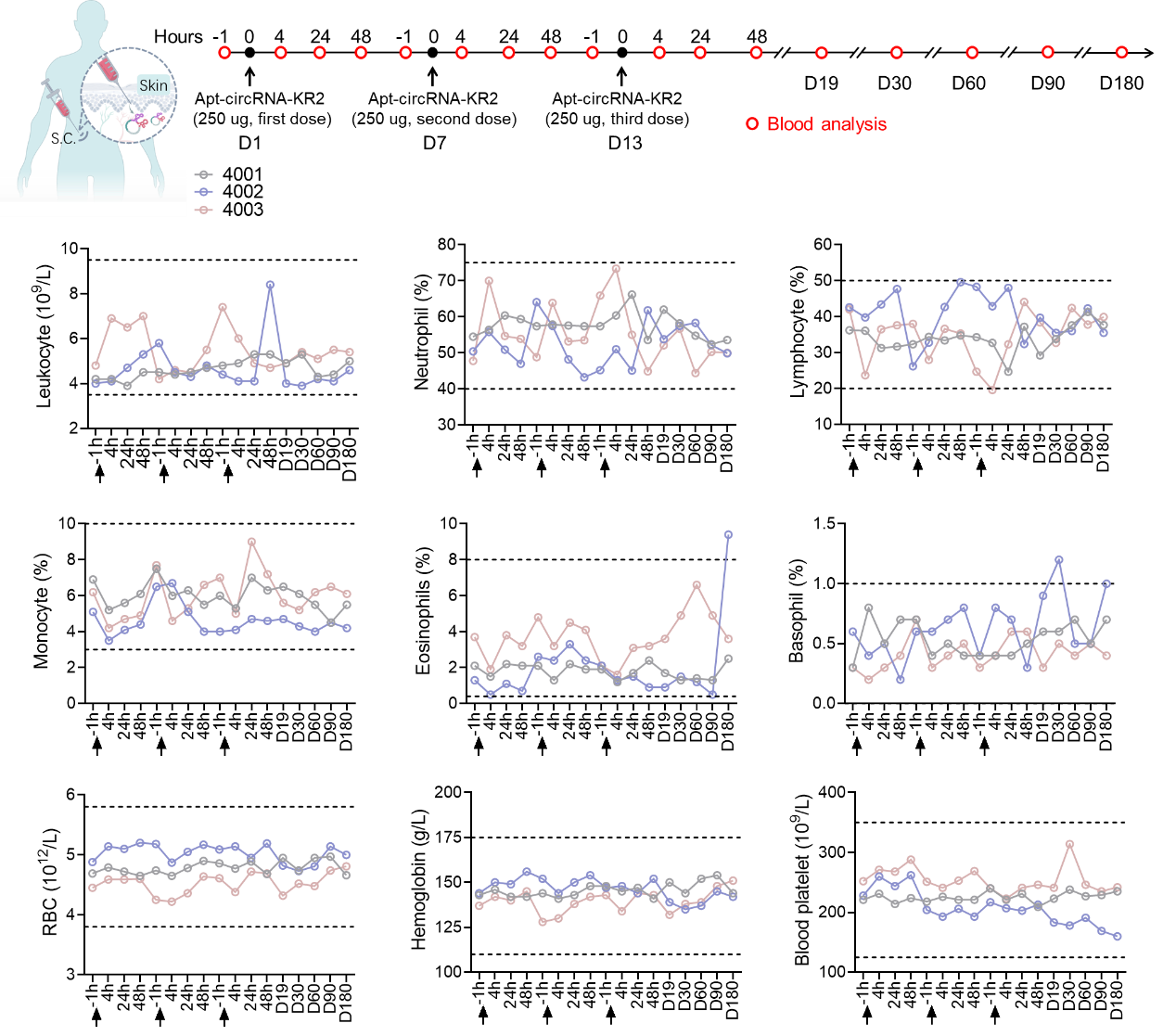

**Fig. S22. Tolerability of Apt-circRNA vaccine in human.** Hematologic parameters of 3 healthy volunteers after immunization with three doses of Apt-circRNA-KR2 vaccine (250 μg; n=3). Black dashed line represents the range of normal individuals. Data represent mean ± s.e.m.

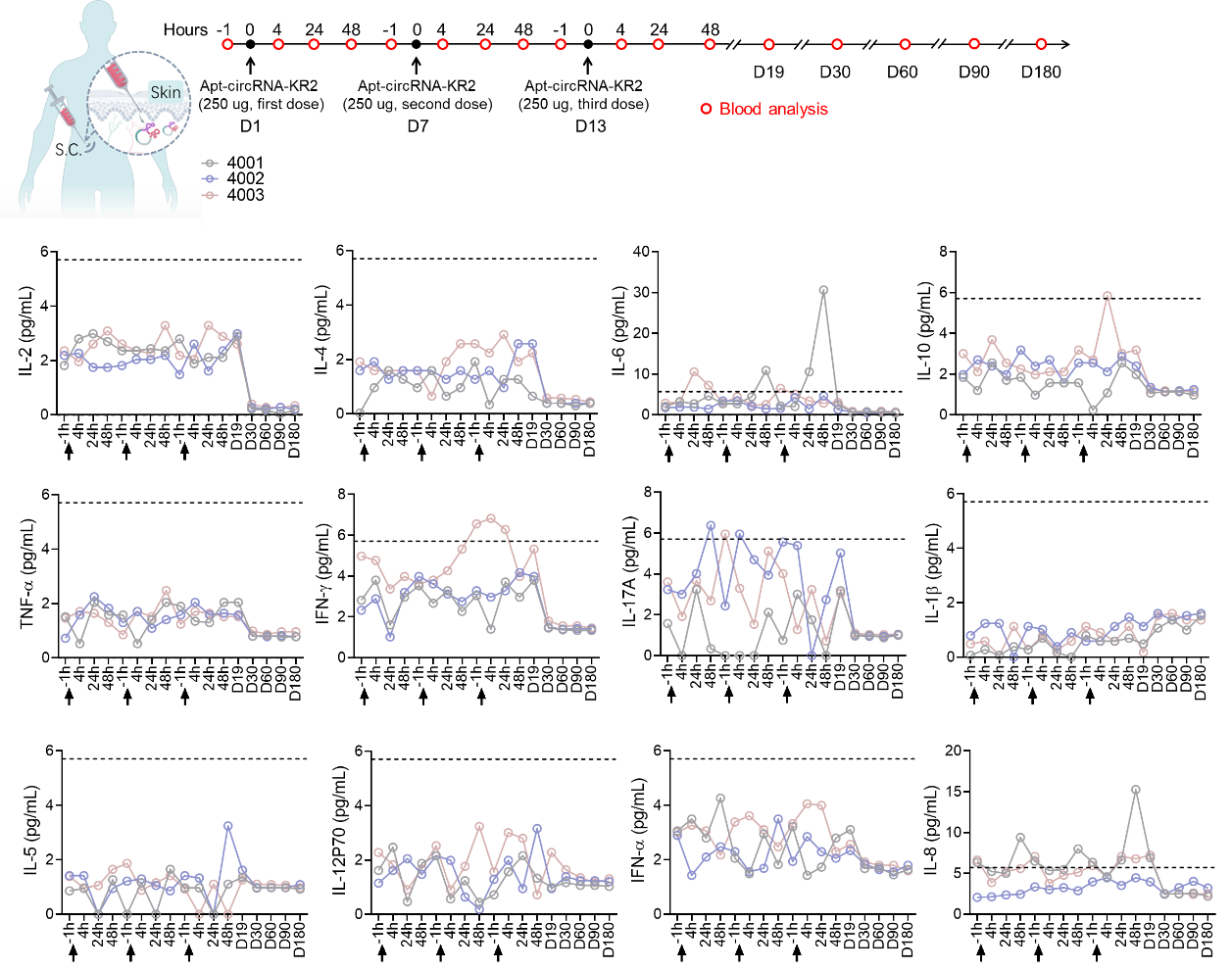

**Fig. S23. Tolerability of Apt-circRNA vaccine in human.** Serum cytokines of 3 healthy volunteers after immunization with three doses of Apt-circRNA-KR2 vaccine (250 μg; n=3). Black dashed line represents the range of normal individuals. Data represent mean ± s.e.m.

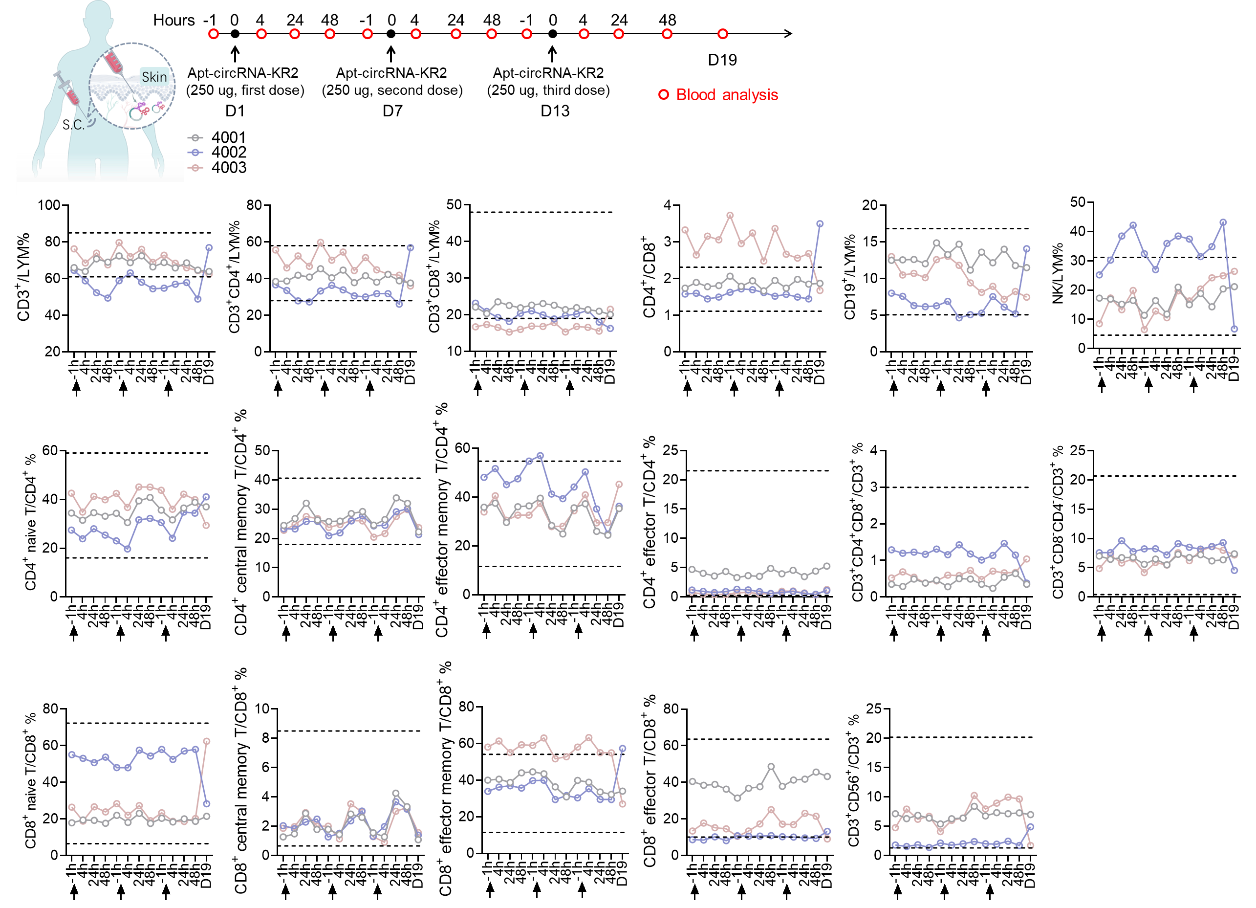

**Fig. S24. Tolerability of Apt-circRNA vaccine in human.** The immune cell subsets of 3 healthy volunteers after immunization with three doses of Apt-circRNA-KR2 vaccine (250 μg; n=3). Black dashed line represents the range of normal individuals. Data represent mean ± s.e.m.

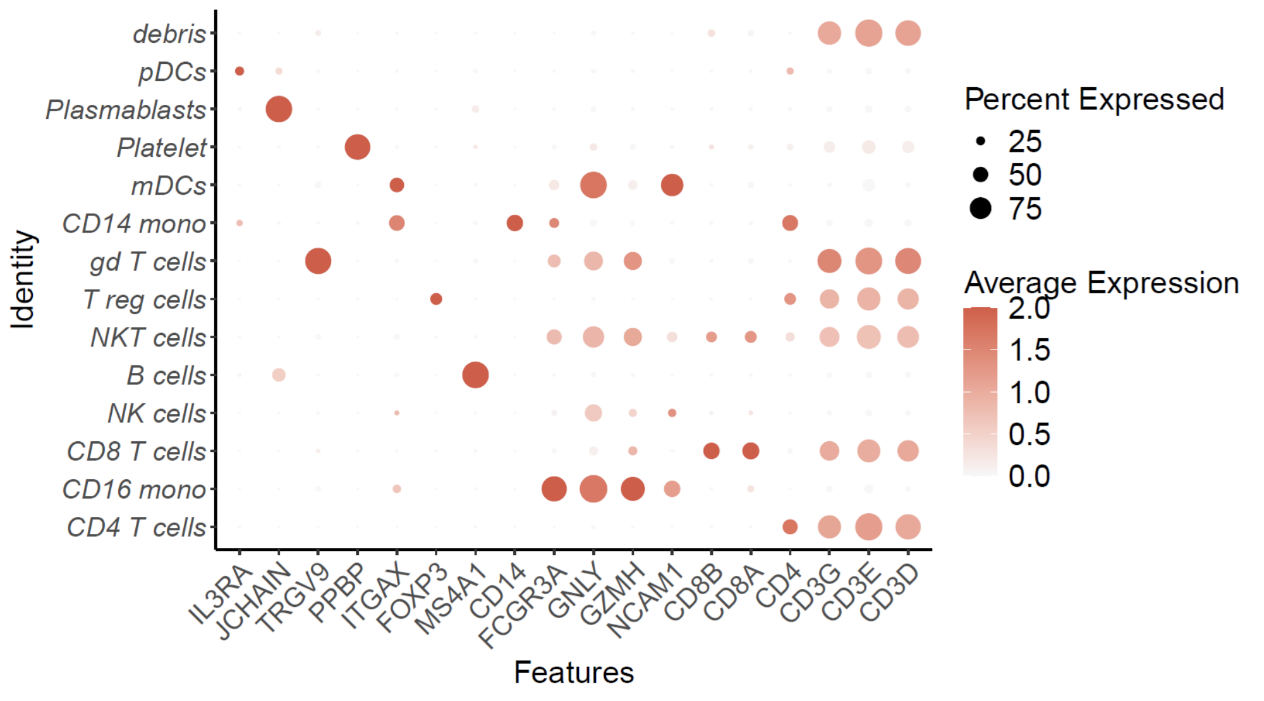

**Fig. S25. Dot plot depicting cell type marker genes across 14 distinct cell types from human PBMC samples.**

**Table S1. List of antibodies used in this study.**

| Targets | Fluorochromes | Vendors | Catalogue # |
| --- | --- | --- | --- |
| CD45 | Brilliant Violet 421 | BioLegend | 103133 |
| CD11c | PerCP-Cy5.5 | BD | 560584 |
| CD11c | PE/Cy7 | BD | 561022 |
| CD11b | BV650 | BioLegend | 101239 |
| CD8a | AF647 | BioLegend | 100727 |
| CD4 | PerCP/Cy5.5 | BioLegend | 100433 |
| CD80 | FITC | BioLegend | 104705 |
| CD86 | APC | BioLegend | 105011 |
| I-A/I-E | APC | BioLegend | 107613 |
| I-A/I-E | PE | BD | 562010 |
| F4/80 | PE | BioLegend | 111603 |
| F4/80 | APC/Cyanine7 | BioLegend | 123117 |
| NK1.1 | APC/Cyanine7 | BioLegend | 108723 |
| NK1.1 | BV 605 | BioLegend | 108753 |
| CD3 | PerCP/Cy5.5 | BD | 561108 |
| CD279(PD-1) | Brilliant Violet 421 | BioLegend | 135217 |
| IFN-γ | PE | BioLegend | 505807 |
| TNF-α | APC/Cyanine7 | BioLegend | 506343 |
| CD44 | PE/Cy5 | BioLegend | 103009 |
| CD62L | FITC | BioLegend | 104405 |
| CD25 | FITC | BioLegend | 101907 |
| FoxP3 | Alexa Fluor 647 | BioLegend | 126407 |
| CD279(PD-1) | N/A | Bio X Cell | CP162 |
| CD152(CTLA-4) | N/A | Bio X Cell | CP146 |
| CD4 | N/A | Bio X Cell | BP0003-3 |
| CD8α | N/A | Bio X Cell | BP0117 |
| CD20 | N/A | Bio X Cell | BE0356 |
| CD45R/B220 | FITC | BioLegend | 103205 |
| EIF2AK2/PKR | N/A | Thermo Fisher | 18244-1-AP |
| FLAG M2 | N/A | Sigma-Aldrich | F1804 |
| $\boldsymbol{\beta}$-actin | N/A | Thermo Fisher | MA1-140 |
| CD45RA | FITC | Beckman Coulter | A07786 |
| CD197 | PE | Beckman Coulter | B30632 |
| CD19 | PC5 | Beckman Coulter | A07771 |
| CD56 | PC7 | Beckman Coulter | A21692 |
| CD4 | APC | Beckman Coulter | IM2468 |
| CD8 | APC700 | Beckman Coulter | C41196 |
| CD3 | APC750 | Beckman Coulter | C41176 |
| CD45 | KRO | Beckman Coulter | C41157 |
| Cytokine detection kit | N/A | Seager | P110100403 |

**Table S2. Peptide sequences.**

| **Peptide names** | **Sequences** |
| --- | --- |
| SIINFEKL | SIINFEKL |
| FLAG | DYKDDDDK |
| Adpgk | ASMTNMELM |
| HPV-16 E7_43-62_ | GQAEPDRAHYNIVTFCCKCD |
| Trp2_177-190_ | ANCSVYDFFVWLHY |
| gp100_23-33_ | ALKVPRNQDWL |
| KRAS (2-25) G12V | TEYKLVVVGAVGVGKSALTIQLIQ |
| KRAS (2-25) G12D | TEYKLVVVGADGVGKSALTIQLIQ |
